# Mechanoreceptor signal convergence and transformation in the dorsal horn flexibly shape a diversity of outputs to the brain

**DOI:** 10.1101/2022.05.20.492853

**Authors:** Anda M. Chirila, Genelle Rankin, Shih-Yi Tseng, Alan J. Emanuel, Carmine L. Chavez-Martinez, Dawei Zhang, Christopher D. Harvey, David D. Ginty

## Abstract

The encoding of touch in the spinal cord dorsal horn (DH) and its influence on tactile representations in the brain are poorly understood. Using a range of mechanical stimuli applied to the skin, large scale *in vivo* electrophysiological recordings, and genetic manipulations, here we show that neurons in the mouse spinal cord DH receive convergent inputs from both low- and high-threshold mechanoreceptor subtypes and exhibit one of six functionally distinct mechanical response profiles. Genetic disruption of DH feedforward or feedback inhibitory motifs, comprised of interneurons with distinct mechanical response profiles, revealed an extensively interconnected DH network that enables dynamic, flexible tuning of postsynaptic dorsal column (PSDC) output neurons and dictates how neurons in primary somatosensory cortex respond to touch. Thus, mechanoreceptor subtype convergence and nonlinear transformations at the earliest stage of the somatosensory hierarchy shape how touch of the skin is represented in the brain.

## Introduction

Our ability to perceive and react to mechanical forces acting on our skin is mediated by the sense of touch. Touch perception begins with activation of physiologically distinct classes of cutaneous mechanosensory neurons and propagation of their signals to the spinal cord dorsal horn (DH) and brainstem. Fundamental to understanding the neurobiological basis of touch perception is knowing how signals from primary mechanosensory neurons are represented and transformed as they ascend the somatosensory neuraxis to encode salient features of tactile stimuli.

The mechanosensory neurons associated with glabrous (non-hairy) skin include fast conducting Aβ rapidly adapting low-threshold mechanoreceptors (LTMRs) that innervate either Meissner or Pacinian corpuscles (Aβ RA1- and Aβ RA2-LTMRs, respectively), Aβ slowly adapting (SA)- LTMRs that associate with Merkel cells, and A- and C-fiber high-threshold mechanoreceptor (HTMR) subtypes that form free nerve endings terminating in the dermis and epidermis (Arcourt et al., 2017; Bessou and Perl, 1969; Handler and Ginty, 2021; Hill and Bautista, 2020; Johnson, 2001; Perl, 1968). Aβ-LTMRs exhibit predominantly two response profiles to sustained skin indentations: Aβ RA-LTMRs fire action potentials at the stimulus onset and offset, while Aβ SA-LTMRs produce sustained spiking patterns and lack an OFF response (Handler and Ginty, 2021; Horch et al., 1977; Johnson, 2001; Owens and Lumpkin, 2014). Moreover, while all Aβ LTMR subtypes can entrain to mechanical vibrations, the force thresholds for activation of both Meissner and Pacinian Aβ RA-LTMRs are uniquely vibration frequency dependent: Meissner corpuscle Aβ RA1-LTMRs optimally encode mechanical vibrations in the ‘flutter’ vibration range (40-100Hz) while Pacinian corpuscle Aβ RA2-LTMRs encode high-frequency vibrations, ranging from 100 to over 500Hz (Freeman and Johnson, 1982; Handler and Ginty, 2021; Johnson, 2001; Mountcastle et al., 1967). In comparison, A- and C-fiber HTMRs require high forces for activation and are typically slowly adapting, firing during initial skin contact and repetitively during sustained contact at high mechanical forces (Arcourt *et al*., 2017; Burgess and Perl, 1967; Cain et al., 2001; Dubin and Patapoutian, 2010; Koltzenburg et al., 1997; Warwick et al., 2021). LTMR and HTMR subtypes also differ in their receptive field (RF) properties. In addition, while the RFs of individual mechanosensory neuron populations homotypically tile the skin, different classes of mechanoreceptors exhibit extensive heterotypic overlap (Kuehn et al., 2019; Neubarth et al., 2020). Thus, tactile stimuli can activate distinct combinations of mechanoreceptor subtypes to produce unique ensembles of impulses propagating from the skin to engage the central touch circuitry.

How are signals from distinct mechanoreceptor subtypes combined in the CNS to generate central representations of touch? The classic view has held that segregated Aβ LTMR signals are conveyed separately via dedicated ascending pathways to drive distinct neuronal populations in somatosensory cortex and generate discriminative touch percepts of form and texture, object orientation, direction and speed of motion, and vibration (Johnson, 2001). Mechanosensory pathways that ascend from the spinal cord to the brain include: 1) the “direct dorsal column pathway”, which carries Aβ LTMR signals directly to the brainstem dorsal column nuclei (DCN) where second order neurons project to the somatosensory thalamus and from there to primary somatosensory cortex (S1); 2) the “indirect dorsal column pathway”, which conveys mechanosensory signals from the spinal cord to the DCN via post-synaptic dorsal column (PSDC) output neurons; and 3) the anterolateral pathway, which transmits signals from the spinal cord directly to the somatosensory thalamus, lateral parabrachial nucleus, and other brain regions that process signals for nociceptive and affective aspects of somatosensation.

Contrary to the classic view of segregated mechanosensory modalities, recent studies indicate that tactile responses in S1 reflect subcortical convergence of Aβ LTMR subtype signals (Emanuel et al., 2021; Pei et al., 2009; Suresh et al., 2021). Evidence for Aβ LTMR signal integration has also been observed in the DCN and somatosensory thalamus (Emanuel et al., 2021; Suresh et al., 2021). Neurons in the macaque DCN, for example, exhibit spatially complex RFs, intermediate rates of adaptation to sustained skin indentation, and orientation tuning, which likely reflect mixing of Aβ LTMR subtype inputs (Suresh et al., 2021). Thus, the emerging view is that signals from physiologically diverse mechanosensory neurons converge early in the somatosensory hierarchy to generate complex tactile feature representations. Indeed, the spinal cord DH may serve as an initial locus of mechanoreceptor subtype signal integration and complex tactile feature representations. In support of this idea, the vast majority of Aβ LTMR synapses are localized to the DH, with relatively few residing in the DCN (Bai et al., 2015; Brown, 1968; Brown et al., 1980; Brown et al., 1981; Brown et al., 1978; Brown et al., 1991; Perl et al., 1962; Petit and Burgess, 1968). Moreover, most if not all other cutaneous somatosensory neuron types, including hairy skin-innervating C-LTMRs, Aδ-LTMRs, and HTMRs, synapse exclusively in the DH (Bai et al., 2015; Kuehn et al., 2019; Li et al., 2011; Olson et al., 2017; Seal et al., 2009; Zylka et al., 2005). Thus, the vast majority of mechanoreceptor synapses reside within the DH. In addition, recent anatomical and *in vitro* electrophysiological analyses point to crosstalk of peripheral mechanosensory channels within the DH (Abraira et al., 2017; Kuehn et al., 2019; Li et al., 2011). Moreover, classic electrophysiological studies in the cat indicate that output neurons of the deep dorsal horn, the PSDC projection neurons, are exquisitely mechanically sensitive, have large, complex RFs, and can exhibit RF plasticity (Brown et al., 1983; Brown and Fyffe, 1981; Brown et al., 1986; Noble and Riddell, 1988; 1989). It is also noteworthy that the DH is predominantly comprised of locally projecting, morphologically and physiologically diverse inhibitory and excitatory interneuron types primed to support sensory input computations, with only a relatively small number of output neurons (Abraira and Ginty, 2013; Abraira et al., 2017; Choi et al., 2020; Gatto et al., 2021; Koch et al., 2017; Moehring et al., 2018; Paixao et al., 2019). Despite this, the *in vivo* response properties of DH neuron types, the nature and extent of mechanoreceptor subtype convergence within the DH, and the contributions of DH mechanosensory coding to tactile representations at higher levels of the somatosensory hierarchy are largely unknown.

Here, we use a range of mechanical stimuli, large scale *in vivo* electrophysiological recordings in conjunction with unsupervised clustering, and mouse genetic manipulations to ask how cutaneous tactile stimuli are encoded by DH interneurons and PSDC output neurons. Our findings indicate that the DH has a highly interconnected network architecture that receives extensively convergent LTMR and HTMR signals and transforms them into a diverse range of PSDC output signals. Thus, the DH flexibly shapes PSDC output signals that ascend via the indirect dorsal column pathway to the DCN where they combine with unmodified Aβ LTMR signals of the direct dorsal column pathway to dictate how touch is represented in the brain.

## Results

### Functional diversity of mechanosensory responses in a large sampling of dorsal horn neurons

As an inroad to understanding the neuron types that mediate somatosensory processing in the spinal cord DH, a number of studies have begun to define its cellular and synaptic architecture using molecular profiling and *in vitro* electrophysiological and morphological approaches (Abraira et al., 2017; Bourane et al., 2015; Choi et al., 2020; Graham and Hughes, 2020; Grudt and Perl, 2002; Haring et al., 2018; Koch et al., 2017; Lima et al., 1993; Lu and Perl, 2005; Osseward et al., 2021; Polgar et al., 1999; Rosenberg et al., 2018; Russ et al., 2021; Sathyamurthy et al., 2018). We sought to complement and extend these prior analyses using *in vivo* electrophysiology to address *(1)* the *in vivo* mechanical response properties of DH neurons, including genetically defined interneurons and tagged PSDC output neurons, *(2)* the contributions of LTMR subtypes, HTMRs, and local circuit dynamics in shaping physiological responses across the DH interneuron network and PSDC output neurons, and *(3)* the contributions of DH mechanosensory information processing to responses observed in S1 and to somatosensory behaviors. We first developed a preparation for *in vivo* spinal cord electrophysiology that allowed us to record simultaneously from dozens of lumbar DH neurons while delivering well-controlled static and dynamic mechanical stimuli to the plantar surface of the hindpaw (Figure 1B). Step indentations spanning the mechanical activation thresholds of both LTMRs and HTMRs (1mN-75 mN) were delivered to paw glabrous skin to assess force intensity tuning of DH neurons residing within spinal cord lamina I through lamina V and allow for direct comparisons to responses of primary mechanoreceptors as well as neurons at higher stages of the somatosensory hierarchy (Figures 1A, S1C). Indentations within this force range were considered innocuous because they did not evoke a nociceptive behavioral response in awake mice (Emanuel et al., 2021). For spatial RF measurements, the same step indentations were applied to different locations across the paw. In addition, vibratory stimuli of varying force amplitudes (1-40mN) and frequencies (2-120 Hz) were delivered to the RF center. A total of 5060 neurons from 142 mice were recorded in the initial stages of this analysis (Figure S1D).

**Figure 1.**
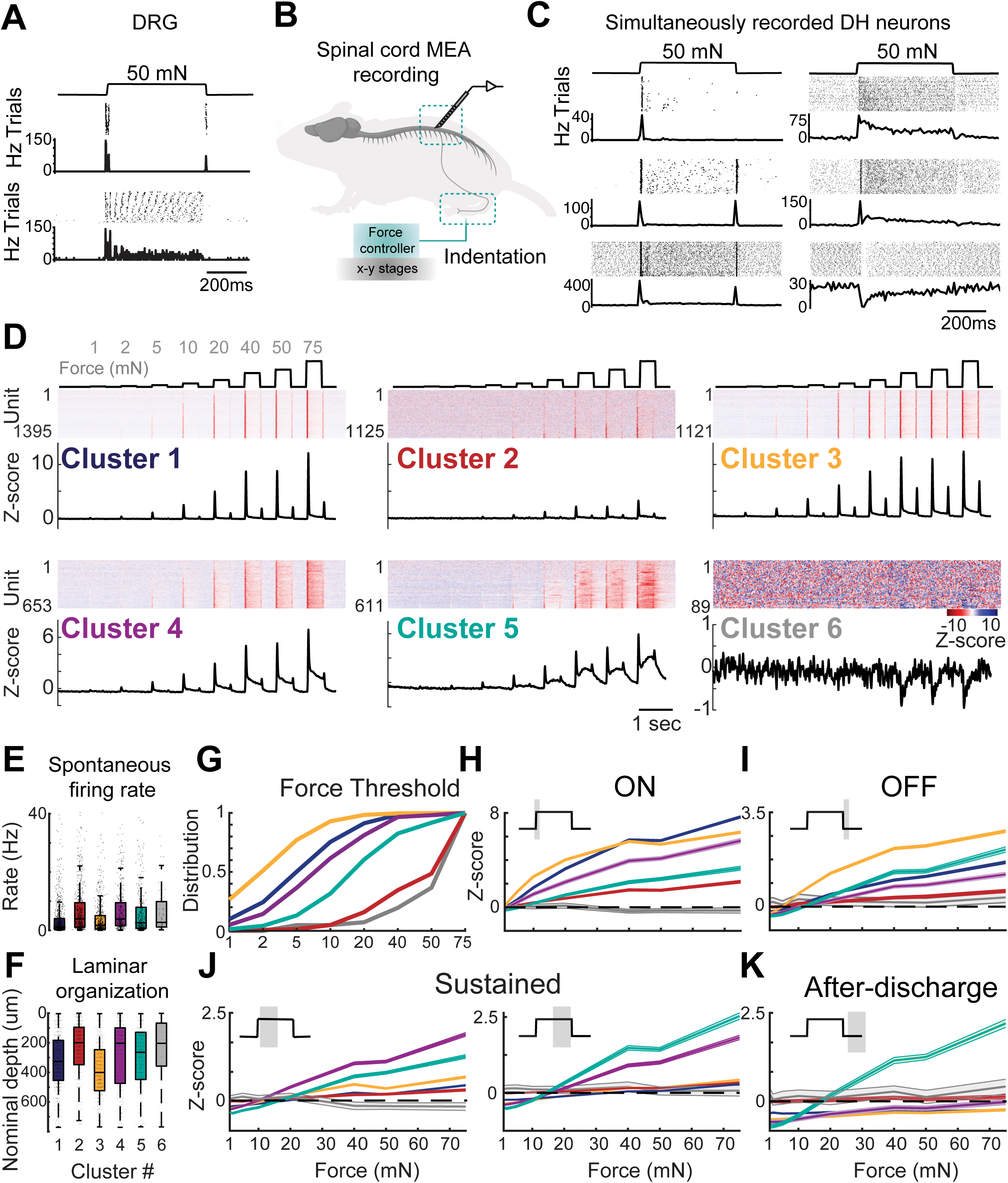
Functional diversity of mechanosensory responses in DH neurons. A. *In vivo* MEA recordings from L4 DRG neurons in anaesthetized mice. Spike raster plots and peristimulus time histograms (PSTHs) aligned to 50-mN indentation step from example Aβ RA-LTMR (top) and Aβ SA-LTMR (bottom). PSTHs are 10ms bins. B. Experimental configuration. L4 spinal cord MEA recordings were performed using 32- channel linear silicon probes in anaesthetized mice while delivering a set of standardized mechanical stimuli to the ventral hindpaw using a force controller atop a X-Y motorized stage. C. Example raster plots and PSTHs from simultaneously recorded dorsal horn neurons during 50-mN step indentations. Top, spike raster; bottom, average PSTH. Histograms are in 10ms bins. D. Z-scored firing rates for all units corresponding to 6 principal functional classes as revealed by unbiased k-medoid clustering (Figure S1). Top, force traces ranging in intensity from 1 to 75- mN aligned to the heatmaps of Z-scored firing rates for each functional group. Functional groups are arranged based on relative abundance, and units in each functional class are sorted by their response thresholds to step indentations. Number of units in each functional group is indicated on the y-axis. Bottom, average PSTHs for each functional group. E. Summary boxplots of spontaneous firing rates for DH units in each functional group (median ± i.q.r. spontaneous firing rates for each functional group: 1.1±3.9Hz; 4.1±8.2Hz; 1.7±4.3Hz; 2.6±7.2Hz; 3.9±8.3Hz; 4.0±12.2Hz. Median ± i.q.r. laminar distribution for each functional group: 341.5±280.2 µm; 198.7±252.0 µm; 400.4±275.5 µm; 294.5±338.6 µm; 220.4± 351.1 µm; 214.0± 280.3 µm. Boxplots: median (line), quartiles (box), minimum and maximum (whiskers). F. Summary boxplots of laminar location for DH units in each functional group. G. Cumulative distribution analysis of mechanical force thresholds for each functional group. **H-K.** Tuning curves i.e. mean (±SEM) firing rates for DH units in each functional group responding to 1mN-75mM step indentations at the onset (ON, 10-50 ms after step onset), offset (OFF; 10-50ms after step offset); sustained (50-250 and 250-500 ms after step offset, respectively) and after-discharge (50-250 ms after step offset) periods. See Table S1 for statistical analyses.

Remarkably, the vast majority of lumbar DH neurons recorded (92%) responded to 500 ms innocuous force steps applied to the paw, and these neurons displayed a broad range of functional response profiles (Figure 1C). To assess the extent of this functional diversity, we used an unsupervised clustering procedure to compare response profiles across all recorded units (Figure S1D). We first used principal component analysis (PCA) to extract features of indentation-evoked signals and then performed k-medoid clustering (Figure S1D, Methods). This analysis divided DH units into six principal functional groups that differ in their responses across the full range of indentation intensities (Figure 1D; arranged based on relative abundance). A similar fraction of broad response types was consistent across experiments (Figure S1E).

DH neurons across the six functional clusters differed in their sensitivity to step indentations, with mechanical activation thresholds and responses during the sustained phase of step indentations tiling innocuous force space (Figure 1G, J). DH neurons in cluster 3 were the most sensitive, with 30% of these neurons responding to 1mN indentation steps, the lowest force amplitude tested (Figure 1D, G). By contrast, neurons in functional cluster 2 and 6 were the least sensitive, with neurons in cluster 2 exhibiting minimal responses at lower forces and less plateauing at higher forces. Cluster 6 harbors neurons uniquely inhibited by high intensity force steps. DH functional clusters were also distinguished by their spiking patterns at the onset (ON), offset (OFF), and during the sustained portion of step indentations (Figure 1D, H-K). Units in functional clusters 1 and 2 exhibited transient response profiles, with robust spiking during the ON phase and little to no response during the sustained portion of the step (Figure 1H-J). Cluster 1 is notable in that neurons in this group often lacked an OFF response (Figure 1D, I). Neurons in clusters 3, 4 and 5, on the other hand, exhibited prominent sustained responses that were most pronounced at higher forces (Figure 1J). DH functional types also differed in their spontaneous firing rates (Figure 1E), with neurons in functional clusters 2, 4, 5 and 6 having on average higher spontaneous firing rates than neurons in clusters 1 and 3. Surprisingly, neurons within different functional groups were broadly distributed across laminar depth (Figure 1F), indicating that low-threshold mechanosensory information is not confined to the LTMR-recipient zone (LTMR-RZ), as had been suggested from histological analysis of LTMR terminals (Abraira et al., 2017). It is noted, however, that units in the least sensitive clusters 2 and 6 tended to reside more superficially and units in the most sensitive cluster 3 tended to reside deeper in the DH (Figure 1F).

We also mapped the excitatory cutaneous RFs of the majority of mechanically sensitive DH units in the dataset. While most DH neurons have spatially continuous RFs (Figure 2A), a subset have spatially discontinuous RFs with multiple hotspots. Regardless of geometry, responses observed in the RF center (hotspot) and surround regions generally differed, with units exhibiting higher sensitivity and spike rates at the RF center measured during the ON, sustained, and OFF portions of step indentations, compared to the RF periphery (Figure S2C, D). We found that RF areas varied across DH functional response types and were exclusively excitatory with no inhibitory surround. Generally, the most sensitive DH neurons tended to have the largest receptive fields.

**Figure 2.**
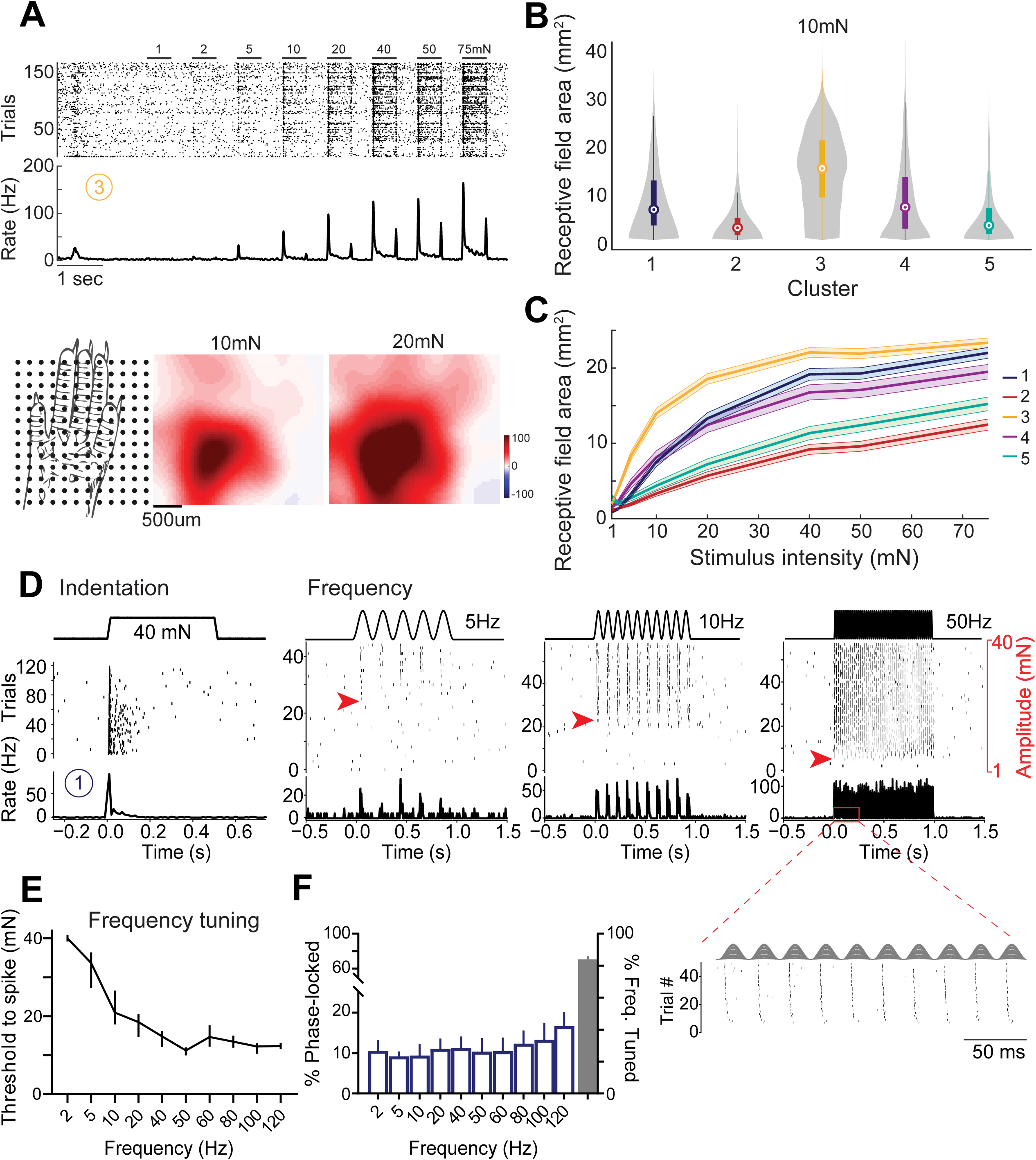
Receptive fields and vibration tuning of DH neuron functional types. **A.** DH linear receptive field organization. Top, spike raster and PSTH from a representative DH unit assigned to functional cluster 3. Each trial shows responses from indentation steps applied to a different randomized location across the paw. Bottom, stimulus locations superimposed on a hindpaw schematic (left), and RF (of the same DH neuron shown above) computed at the onset of 10 and 20mN step indentations (right; see Methods). **B.** Excitatory RF areas at 10mN indentation steps for units across five principal functional groups (number of units and median RF for each functional group: 737units; 6.24 mm^2^; 605 units; 2.34 mm^2^; 1106 units; 14.25 mm^2^; 626 units; 5.5 mm^2^; 488 units; 3.1 mm^2^). P < 0.0001, Kruskal-Wallis H test. **C.** Average receptive field areas vs. indentation depth. Shaded areas: SEM. **D.** Example DH neuron that exhibits both phase-locking and vibration tuning. This example DH unit was assigned to functional cluster 1 based on its indentation response (left), and responses to mechanical vibrations. Note that the force activation threshold for this unit decreases with increasing frequency of vibration, indicative of frequency tuning (red arrows). Inset shows spikes phase-locked to the periodic cycle of the 50Hz vibration. **E.** Frequency tuning curves (mechanical activation threshold vs. stimulation frequency) for all DH neurons in the dataset. Data represent median ± 95%CI. **F.** Left: Percentage of DH neurons that were phase-locked across all vibratory frequencies tested; Right: Percentage of frequency-tuned DH neurons (83.9±2.3 %). Data represent mean ± SEM. See Table S1 for statistical analyses.

Indeed, while units in cluster 3, the most sensitive cluster, exhibited the largest RFs at 10mN (median 14.75mm^2^), units in cluster 2, which have the highest activation thresholds, exhibited the smallest RFs (median 2.34 mm^2^) (Figure 2B, C). Given the low indentation forces that activate Aβ RA- and Aβ SA-LTMRs (median 5mN; Emanuel et al., 2021), their small localized receptive fields, and their extensive homotypic tiling across glabrous skin, this analysis indicates that RFs for virtually all DH units in the dataset and representing all excitatory functional clusters are considerably larger than the RFs measured for individual Aβ-LTMRs, suggesting a high degree of convergence of primary mechanosensory neuron signals onto individual DH neurons (see below).

We also investigated how neurons across the DH encode vibratory stimuli. The extent of phase-locking to individual phases or cycles of vibratory stimuli as well as vibration tuning, i.e., frequency-dependent sensitivity measured across a range of forces, were assessed for the majority of units in the dataset (Figure 2D-F). Most neurons exhibited little or no phase-locked spike entrainment to the full range of vibration frequencies tested. In fact, only a small subset of DH neurons (15%) phase-locked to vibrations up to 120Hz (Figure 2F). In contrast, the majority of DH neurons across all functional clusters showed broad vibration tuning (84%; Figure 2E).

Thus, the sensitivity of most mechanically evoked responses in DH neurons increased (i.e., force thresholds decreased) as the frequency of vibration increased, at least to 120Hz (Figure 2E).

These findings suggest that most DH neurons receive convergent input, either directly or indirectly, from Meissner corpuscle Aβ RA1-LTMRs and Pacinian corpuscle Aβ RA2-LTMRs, which are most sensitive within the 20-100Hz and 60-500Hz ranges, respectively (Johnson, 2001).

*Genetically defined DH neuron subtypes map onto functionally defined clusters.* We next asked whether DH neurons within particular functional clusters align with known genetically defined DH interneuron populations. We first tested whether DH units across different functional clusters have unique extracellular waveform features that could be predictive of previously defined cell types using K-means clustering on our comprehensive DH waveform dataset (Figure S2 E-G, Methods). While this strategy yielded well-separated clusters with fast-spiking and regular-spiking features on a spike waveform dataset of somatosensory cortex neurons, as previously described (Jia et al., 2019; Niell and Stryker, 2010), spike waveforms from individual DH units were not clearly separable (Figure S3A-C). Therefore, we next combined *in vivo* MEA recordings with optical tagging of genetically defined DH interneuron types, using DH interneuron specific Cre driver lines (Figure 3A), to selectively record from *(1)* broad DH neuron populations subdivided based on neurotransmitter identity and *(2)* a sampling of previously described morphologically and physiologically distinct DH interneuron subtypes that fall within excitatory and inhibitory interneuron subdivisions. The *in vivo* response properties and RFs of genetically labeled DH interneuron populations were then compared to those of DH functional types defined by unbiased clustering.

**Figure 3.**
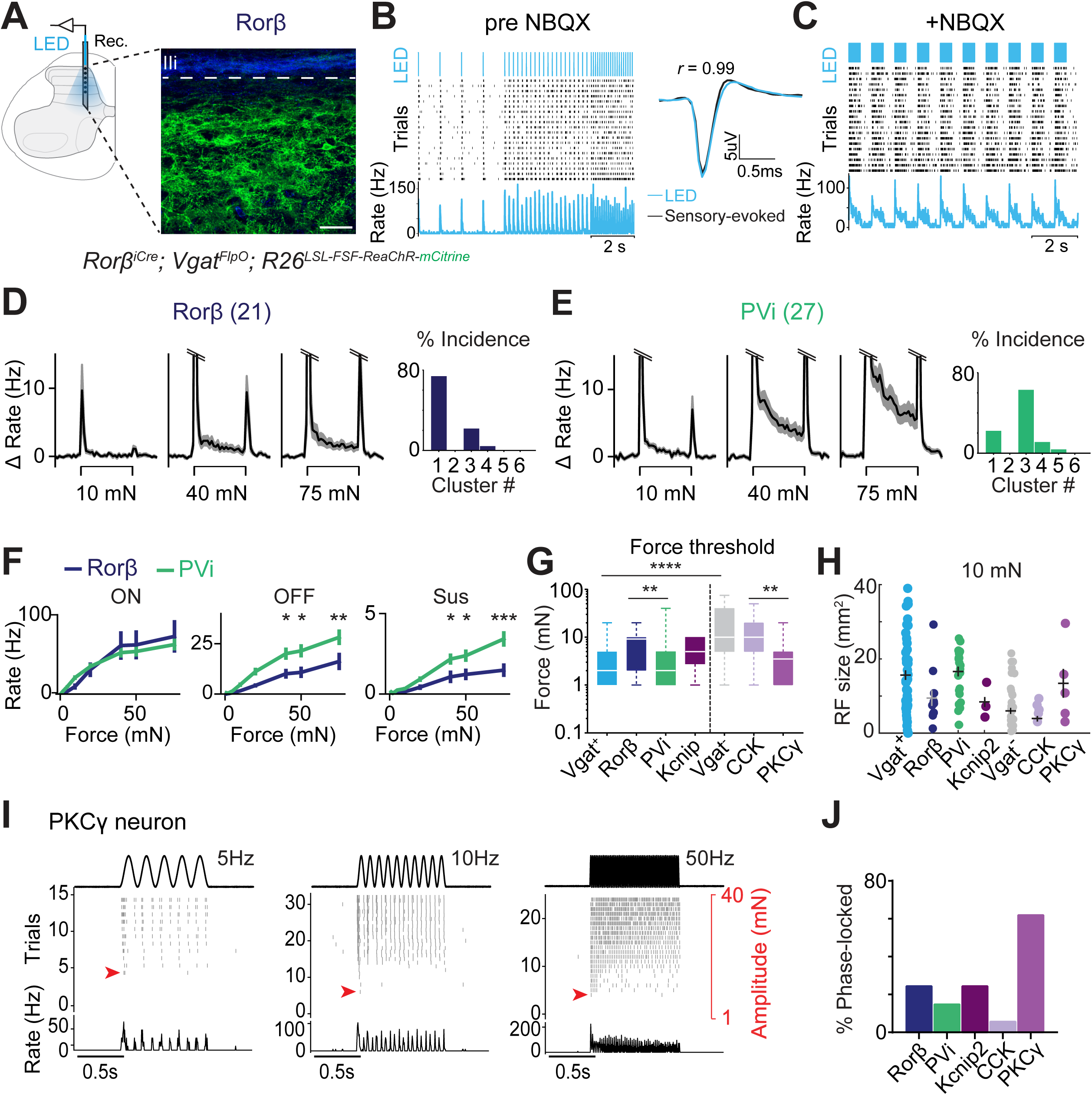
Genetically defined interneurons map onto DH neuron functional types. **A.** Left: *in vivo* experimental set-up and recording configuration for optotagging genetically defined neuron types. A light-emitting diode (LED)-coupled optic fiber was used to optically excite DH neurons expressing an opsin while recording DH units. Right: Sagittal spinal cord section at the recording site shown in the schematic (left) from a *Rorβ^iCre^; Vgat^FlpO^; R26^LSL-FSF-ReaChR-mCitrine^*mouse (green, ReaChR; IB_4_ binding in blue labels lamina IIi). **B.** Optotagging protocol (top) with corresponding light-evoked spike raster and PSTH for an example optotagged Rorβ unit. Right, average indentation- and light-evoked waveforms. Optically tagged neurons display high waveform correlation between light-evoked and sensory- spikes (r=0.99). **C.** NBQX (5mM) was applied to the surface of the cord to block excitatory synaptic transmission at the end of each recording. **H.** Mean indentation-evoked response profiles (firing rate ± SEM) of optotagged Rorβ neurons (n=21 units; 5 mice). The ON response is truncated at 15Hz. Right: Cluster assignment for tagged Rorβ neurons based on their response profiles to static indentation. **I.** As in (H), but for tagged PV_i_ units. PV_i_ neurons have more pronounced sustained responses and are mostly assigned to functional cluster 3 (*PV^2aCre^; Vgat^FlpO^; R26^LSL-FSF-ReaChR-mCitrine^*; n=27; 4 mice). The same optotagging protocol was used for tagging PV_i_ neurons as described for Rorβ neurons, in A-G. **J.** Comparison of the response properties of Rorβ and PV_i_ neurons. Mean indentation evoked firing rates (± SEM) at the onset (ON), offset (OFF), and sustained (Sus) portion of the step indentation for each force. Two-way ANOVA with post hoc Sidak’s test; ON: (F [1, 328] = 0.009453, p = 0.9226); OFF: (F [1, 328] = 22.34, p <0.0001); Sus: (F [1, 328] = 22.14, p <0.0001): ***p < <0.0001, **p < 0.001, *p < 0.05. **K.** Summary of the *in vivo* mechanical force thresholds for all photo-tagged dorsal horn neurons examined in this study. Dashed line separates inhibitory from excitatory interneuron types. Note that inhibitory neurons as a group (Vgat^+^ neurons) are considerably more sensitive than excitatory neurons (Vgat^-^). Boxplots: median (line), quartiles (box), minimum and maximum (whiskers). P <0.0001, Kruskal-Wallis H test. Post hoc Dunn’s test for multiple comparisons. ****p < 0.0001, *p < 0.05. **L.** RF sizes for all optotagged DH neurons in this study computed using 10mN force steps applied across the paw. See also Methods. **M.** Tagged DH neurons exhibit phase-locked responses to vibratory stimuli to varying degrees. Left: example of a phase-locked PKCγ^+^ unit, shown at 3 selected frequencies. **N.** Percentage phase-locked units for optotagged DH interneurons. Note that while most tagged neurons show little or no phase-locked entrainment, a large portion of PKCγ^+^ neurons (62.5%) entrain to vibrations up to 120Hz.

To compare broadly across inhibitory and excitatory interneuron populations, both GABAergic and glycinergic inhibitory interneurons were optotagged using *Vgat^iCre^*; *R26^LSL-ChR2^* mice (Figures S3E, F). This analysis revealed that inhibitory DH interneurons are more sensitive and produced more sustained responses at higher indentation forces than simultaneously recorded untagged units, which are putative excitatory neurons (Figures 3G, S3E, F). Moreover, as a population, inhibitory interneurons exhibited larger receptive fields than their excitatory interneuron counterparts (Figure 3H). Despite these differences, there was substantial heterogeneity among the inhibitory and excitatory DH neuron response profiles, as illustrated by functional cluster assignments of both subtypes (Figures S3E, F) as well as their RF areas (Figure 3H). These findings are consistent with prior *in vitro* recordings that had divided inhibitory and excitatory DH populations into distinct subsets with unique combinations of intrinsic physiological and synaptic properties.

We also sampled five genetically defined DH interneuron populations through optotagging using LTMR-RZ specific Cre-driver lines and either ChR2 or ReaChR actuator mouse lines (Figures 3A-C, S3A-D; see Methods) to assess their *in vivo* responses to the same battery of mechanical stimuli used for the random recordings. This analysis showed that each of the five genetically labeled populations tested has a relatively homogenous mechanical response profile and differentially map predominantly onto one or more of the six functional response profiles from the large dataset (Figure 1D, S3F-J). Rorβ^+^ inhibitory interneurons (optotagged using *Rorβ^iCre^;Vgat-FlpO; R26^LSL-FSF-ReaChR^* mice) exhibited transient responses to skin indentations, with little to no sustained response and a modest OFF response at low forces, and thus the majority of these interneurons mapped onto DH functional cluster 1 (Figure 3D). In contrast, PVi inhibitory interneurons (optotagged using *PV^2aCre^;Vgat-FlpO; R26^LSL-FSF-ReaChR^* mice) exhibited some of the lowest force thresholds in the dataset as well as pronounced sustained and OFF responses to step indentations, and thus these interneurons were mostly assigned to DH functional cluster 3 (Figure 3E). Also consistent with their cluster 3 assignment is the observation that PVi neurons displayed some of the largest RFs in the dataset (Figure 3H).

Another genetically labeled inhibitory population, Kcnip2 interneurons (optotagged using *Kcnip^CreER^; R26^LSL-ChR2^* mice) that form axodendritic synapses within the LTMR-RZ (Abraira et al., 2017), had a distinct cluster assignment, a reflection of their unique sustained firing patterns in response to indentation, high spontaneous firing rates, and small RFs (Figures 3G,H, and S3I, L). In contrast to these three inhibitory interneuron types, the large group of CCK^+^ excitatory lineage neurons exhibited some of the smallest RFs (Figure 3H). Interestingly, optotagged PKCγ^+^ excitatory interneurons, a relatively small subset of spatially restricted DH excitatory interneurons (labeled using *PKC*γ*^CreER^; R26^LSL-ChR2^* or *PKC*γ*^CreER^; Lbx1^FlpO^; R26^LSL-FSF-ReaChR^* mice) were found to be a physiologically homogeneous population with higher sensitivity and larger receptive fields compared to other excitatory interneuron populations (Figures 3G, H, and S3K). Responses of the genetically labeled LTMR-RZ interneurons to mechanical vibrations were also assessed. PKCγ excitatory interneurons stood out in this analysis, as 60% of this population precisely phase-locked to vibratory stimuli at frequencies between 10 and 120 Hz, compared to only 6% of the broad CCK excitatory interneuron population (Figure 3I, J). On the other hand, the majority of units across both the excitatory and inhibitory neuron cohorts and each of the five optotagged interneuron populations exhibited vibration frequency-dependent sensitivity (Figure S3N).

Together, these findings point to a broadly distributed arrangement of mechanically sensitive DH neuron types that fall into six principal functional clusters, likely corresponding to one or more genetically defined interneuron types with distinct intrinsic physiological, morphological, and synaptic properties.

### LTMR and HTMR signal convergence shapes touch-evoked responses in DH neurons

To assess the contributions of mechanoreceptor subtype inputs to the range of *in vivo* physiological response profiles of DH neurons, we next used sensory neuron loss-of-function and gain-of-function manipulations in conjunction with *in vivo* DH MEA recordings. We first asked whether DH neuron responsivity to skin indentation is dependent on the mechanosensory ion channel Piezo2 (Ranade et al., 2014). Indeed, DH neurons of mice in which *Piezo2* was deleted from all neurons below cervical level 2 (*Cdx2-Cre; Piezo2^flox/flox^* mice; Lehnert et al., 2021) displayed virtually no responses to indentation force steps up to 75mN, while pinch-evoked responses were intact (Figure S4A-D). We next assessed the contributions of Aβ RA- and Aβ SA-LTMRs to distinct DH response profiles using genetic ablation strategies that result in loss of Meissner corpuscles and their associated Aβ RA-LTMRs (*Advillin^Cre^; TrkB^flox/flox^* mice referred to as *TrkB^cKO^*(Neubarth et al., 2020) or loss of both Meissner corpuscles/Aβ RA-LTMRs and Merkel cells, which are required for normal Aβ SA-LTMR responses (*Advillin^Cre^; TrkB^flox/flox^; Atoh^flox/flox^* referred to as DKO; Emanuel et al., 2021). We found that DH neurons in mice with disrupted signaling from either Aβ RA-LTMRs alone or both Aβ RA-LTMRs and Aβ SA-LTMRs exhibited dramatically decreased sensitivity compared to control littermates, both at the population level (fraction of units responding at each intensity; Figure 4C, F) and in individual units (response thresholds; Figure 4G). In DKO mice lacking both Aβ RA- and Aβ SA-LTMR signals, ON responses to indentation forces below 20 mN were absent (Figure 4A-F), demonstrating that Meissner corpuscle-associated and Merkel cell-associated Aβ LTMRs mediate light touch responses broadly, across all DH functional response types. DH sensitivity was also diminished, but to a lesser degree, in *TrkB^cKO^* mice that only lack Aβ RA-LTMR signals (Figure 4A-C). In addition, OFF responses to step indentations were absent at low forces and reduced at higher force intensities in both mutants (Figure 4B, E). Phase-locking and vibration sensitivity were also dramatically altered in both *TrkB^cKO^* and DKO mice: DH units from DKO mice were particularly deficient and lacked phase-locked responses at most vibration frequencies tested (Figure 4J). As predicted, a few DH units in DKO mice did exhibit phase-locking at 120Hz (Figure 4H), presumably because high frequency skin vibrations >100Hz at low indentation forces are transduced by Aβ RA2-LTMRs associated with Pacinian corpuscles.

**Figure 4.**
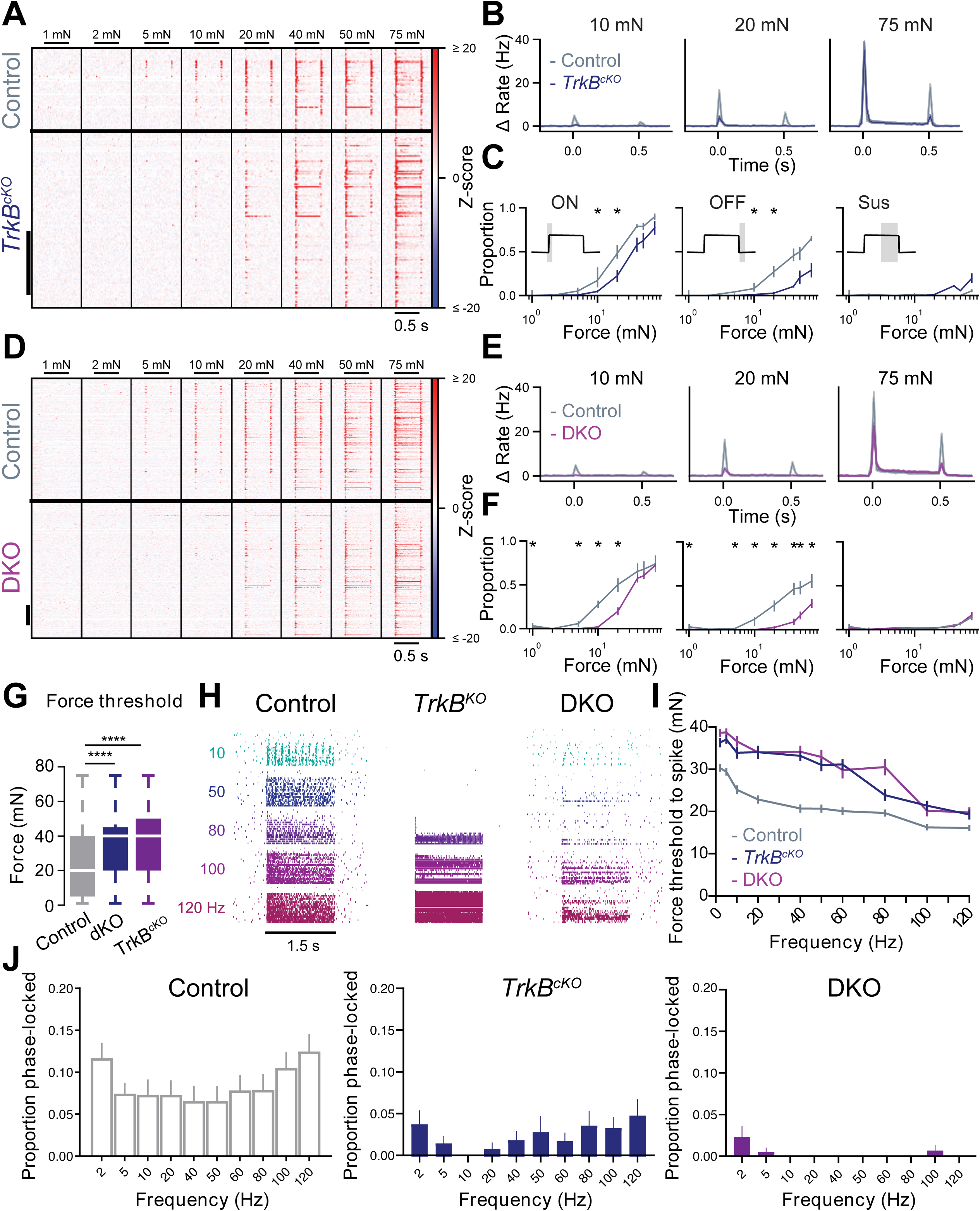
Decreased DH neuron sensitivity in the absence of Aβ-RA-LTMR and Aβ SA- LTMR signals. **A.** Indentation-evoked firing rates for all dorsal horn units from littermate control (*TrkB ^flox/flox^*; top; n=72 units, 3 mice) and *Advillin^Cre^;TrkB^flox/flox^*animals lacking Meissner corpuscles (*TrkB*^cKO^; bottom; n=142 units, 4 mice). Units are sorted by depth. Scale bar: 50 units. **B.** Grand mean baseline-subtracted firing rates (± SEM) to 10, 20, and 75mN step indentations from control (grey) and *TrkB*^cKO^ (dark blue) animals. **C.** Proportion of units from control (grey) and *Advillin^Cre^;TrkB^flox/flox^* animals (dark blue) responding at the onset (left), offset (middle) and sustained (right) phase of step indentation for each force. * p < 0.05 for comparisons between units in *TrkB^flox/flox^* and *TrkB*^cKO^ mice, two-proportions Z test. **D.** As in A, for DH units in littermate controls (top; n=328 units; from 5 recordings in N=4 animals) and DKO (*Advillin^Cre^;TrkB^flox/flox^*; *Atoh1^flox/flox^*; bottom; n=436 units; from 8 recordings in N=4 animals) mice lacking both Meissner corpuscles and Merkel cells. **E.** As in B, for all littermate control (grey) and DKO (purple) DH units. **F.** As in C, for DKO littermate controls (grey) and DKO (purple) units. **G.** Response thresholds from all units in control, *TrkB*^cKO^ and DKO animals. P < 0.0001, Mann-Whitney *U* test. **H.** Example unit responses to mechanical vibrations varying in frequency (from 10 to 120Hz) and amplitude (from 1 to 40mN; ordered by frequency, then amplitude) in control, *TrkB*^cKO,^ and DKO animals. **I.** Frequency- threshold tuning curves in all units from control, *TrkB*^cKO^, and DKO animals. Note the lack of frequency tuning at preferred Meissner frequencies. **J.** Proportion of phase-locked units in control (left), *TrkB*^cKO^ (middle), and DKO (right) animals.

Consistent with this, when force thresholds were calculated at vibration frequencies ranging from 2Hz to 120Hz, both *TrkB^cKO^* and DKO mice lacked frequency tuning below 80Hz but did exhibit normal responsivity in the Pacinian range, above 80Hz (Figure 4I). These findings suggest that signals from Meissner Aβ RA1-LTMRs and Pacinian Aβ RA2-LTMRs converge to generate wide dynamic range vibration frequency-dependent tuning in the majority, and possibly all, DH neurons.

Interestingly, while OFF responses to step indentations, phase-locking to low-frequency vibratory stimuli, and vibration frequency-dependent sensitivity were dramatically compromised in mice lacking Aβ RA-LTMR signals or both Aβ RA- and Aβ SA-LTMR signals, the fraction of DH units spiking during the ON and sustained portions of indentations at forces >20mN was comparable to that of control mice (Figure 4C, F). This finding, along with the observation that the sustained portion of the response to step indentations encodes intensities exceeding those at which Aβ SA-LTMRs saturate, suggests that LTMR and HTMR signals converge onto the majority, and possibly all, mechanically sensitive DH neurons and across cluster types. To further explore this possibility, we generated *Nav1.8^Cre^; Piezo^flox/flox^* mice in which Piezo2 is deleted from a large fraction of small- and medium-diameter neurons, including many HTMRs, but not from Aβ RA- and Aβ SA-LTMRs. As predicted, DH neurons in *Nav1.8^Cre^; Piezo^flox/flox^*mice exhibited no changes in mechanical response thresholds as compared to littermate controls (Figure S4J). However, these mutants did exhibit diminished firing rates at higher indentation forces, above 10mN during the sustained portion of step indentations (Figure S4G-H). This reduction is modest and likely an underestimation of the HTMR contribution to the force-response relationship in the DH due to incomplete deletion of Piezo2 in large diameter CGRP^+^ DRG neurons in *Nav1.8^Cre^; Piezo^flox/flox^* mice (Figure S4E-F).

The loss-of-function analyses were complemented by gain-of-function experiments designed to assess the sufficiency of select LTMR subtypes and HTMRs for activation of DH neurons across different functional cluster types. Testing for sufficiency of individual mechanoreceptors using mechanical stimuli is not feasible because multiple mechanoreceptor subtypes are co-activated, albeit to varying degrees, by mechanical stimulation of the skin. We therefore compared DH neuron responses to mechanical stimuli and responses to optogenetic stimuli applied to the same skin region to selectively activate one or more somatosensory neuron types. Genetic driver lines were used to express ReaChR (Hooks et al., 2015; Lin et al., 2013) in select populations of sensory neurons (see Methods), and brief light pulses were applied to the skin to trigger one action potential in a few sensory neuron (Emanuel et al., 2021). Strikingly, selective optogenetic activation of the cutaneous endings of either Aβ RA-LTMRs or Aβ SA-LTMRs evoked exclusively short-latency responses in the majority of mechanically sensitive DH units (Figure 5B, C, F-H). Thus, even DH neurons with transient mechanical response profiles (e.g., neurons in cluster 1) were driven by optical activation of Aβ SA-LTMRs, which have sustained responses to skin indentation, and DH neurons exhibiting prominent sustained responses (e.g., neurons in clusters 4 and 5) were driven by optical activation of Aβ RA-LTMRs, which have transient responses to skin indentation (Figure 5B, G-H). Furthermore, the spatial RFs of DH neurons mapped with low force mechanical stimuli and patterned optogenetic stimuli to selectively activate Aβ RA-LTMRs or Aβ SA-LTMRs across the skin surface allowed us to compute homotypic convergence estimates for these two LTMR subtypes (Figure 5F). We estimate that, on average, individual DH neurons receive convergent inputs from 10 or more of these Aβ- LTMR subtypes (Figure 5F). These findings, which are consistent with the loss-of-function experiments, indicate an extensive degree of homotypic and heterotypic convergence of Aβ RA-LTMR and Aβ SA-LTMR signals onto most DH neurons and across all cluster types.

**Figure 5.**
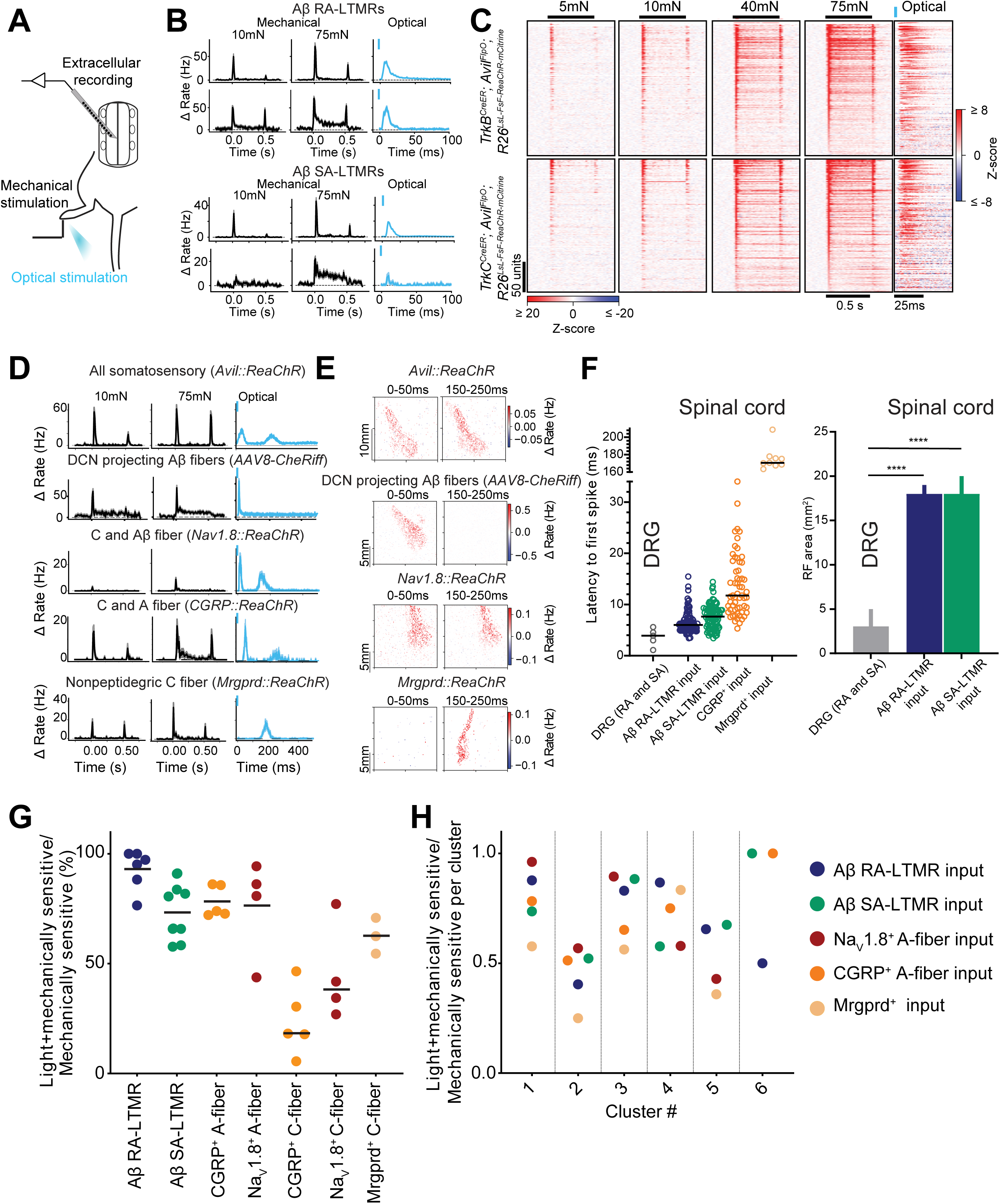
Convergent LTMR and HTMR signals shape DH neuron responses. **A.** Schematic of recording configuration. **B.** Mechanical and optically- evoked PSTHs (mean ± SEM) in example DH units from *TrkB^CreER^*; *Advillin^FlpO^; R26^LSL-FSF-ReaChR-mCitrine^* (used to label Meissner corpuscle associated Aβ RA-LTMRs; top) and from *TrkC^CreER^*; *Advillin^FlpO^; R26^LSL-FSF-ReaChR-mCitrine^* (used to label Merkel cell associated Aβ SA-LTMRs; bottom) mice. Note that selective optical activation of either Aβ RA-LTMRs or Aβ SA-LTMRs evokes firing in DH units with both transient and sustained response profiles to 75mN step indentations. **C.** Z-scored indentation- and optically-evoked responses in all mechanically sensitive (response produced to 75-mN step indentation) DH units. Units are sorted based on the magnitude of ON response. Note the timescale difference between mechanical and optical heatmaps. See Star Methods for optical activation protocol. **D.** As in B, in animals expressing excitatory opsins in all somatosensory (*Advillin^FlpO^; R26^FSF-ReaChR-mCitrine^*); Aβ-DCN projecting Aβ fibers (AAV8-CheRiff DCN injection), Nav1.8^+^ expressing C fibers (*Nav1.8^Cre^; R26^LSL-ReaChR-mCitrine^*); CGRP^+^ HTMRs (*CGRP^CreER^; R26^LSL-ReaChR-mCitrine^*); MRGPRD^+^ nonpeptidergic C fibers (*Mrgprd^CreER^; R26^LSL-ReaChR-mCitrine^*). Note that 500ms ISI was used for optical stimulation to record both A and C fiber inputs to DH neurons. See Start Methods for optical activation protocol. **E.** Optically-evoked RFs revealing both A- and C-fiber RFs in DH neurons. Note somatotopic overlap between A- and C- fiber RFs. **F.** Left: Optically-evoked spike latencies in DRG neurons and spinal cord neurons across different genotypes. Right: Summary optical RF areas in DRG neurons and spinal cord neurons. Bars: median with 95% CI. **G.** Proportion of mechanically sensitive units that respond to optical activation of distinct sensory neuron types in glabrous hindpaw across all genotypes tested. Each marker represents one animal; markers are color-coded by genotype. Bars: mean. **H.** Proportion of mechanically sensitive DH units receiving synaptic input from distinct LTMR and HTMR subtypes per DH functional group. Shaded lines separate adjacent functional clusters. Aβ-RA LTMR input (n=205 units; N=6 mice); Aβ SA-LTMR input (n=215 units; N=8 mice); Nav1.8 input (n=101 units; N=4 mice); CGRP input (n=99 units; N=5 mice); *Mrgprd* input (n=45 units; N=3 mice).

In contrast to Aβ-LTMRs, optogenetic activation of all somatosensory neurons innervating glabrous skin (using *Advillin-FlpO; R26^FSF-ReaChR^* mice) elicited both short and long latency responses in mechanically sensitive DH neurons (Figure 5D). Similar short and long latency responses were observed following electrical stimulation of the skin and were indicative of A-fiber and C-fiber inputs, respectively (Figure S5L). In contrast, skin optogenetic stimulation of DCN-projecting Aβ-fiber sensory neurons (Figure 5D; see Methods) evoked only short latency responses in virtually all mechanically sensitive DH neurons. We also observed that broad optogenetic activation of HTMRs and other sensory neuron types, using both *Nav1.8^Cre^* lineage and *CGRP^CreER^* driver lines, triggered both short (A-fiber) and long (C-fiber; >100ms) latency responses (Figure 5D, F) in the majority of mechanically sensitive DH neurons (Figure 5G, H). In fact, as observed for Aβ RA-LTMRs and Aβ SA-LTMRs, optogenetic activation of fast conducting (A-fiber), CGRP^+^ DRG neurons, which are A-fiber HTMRs (Arcourt et al., 2017; Ghitani et al., 2017; Hoheisel et al., 1994; McCarthy and Lawson, 1990), evoked spiking in the majority of DH neurons and across all functional classes (Figure 5G, H). Also intriguing is the observation that optical RFs for A-fiber and C-fiber evoked responses overlapped somatotopically for the majority of DH units (Figure 5E). Finally, optogenetic activation of Mrgprd^+^ HTMRs evoked exclusively long-latency responses (Figure 5D, F), as expected for these C-fiber neurons, and across DH units corresponding to each of the major functional cluster types (Figure 5H).

Together, these primary sensory neuron loss-of-function and gain-of-function manipulation experiments suggest that parallel signals from physiologically distinct Aβ LTMR subtypes and HTMRs converge extensively in the DH to shape the mechanical responses and RFs of DH neurons across all functional cluster types.

### Sensory-evoked feed-forward and feedback inhibitory circuit motifs exert broad control over DH neuron responses

Our findings point to extensive LTMR subtype and HTMR signal convergence onto the majority of DH interneurons. We next tested the hypothesis that the principal DH interneuron types are functionally inter-dependent, with each type uniquely contributing to mechanical responses and RFs of the others, and ultimately to the signals conveyed by DH output neurons. This was addressed by determining the extent to which two of the genetically defined DH neurons, Rorβ and PVi inhibitory interneurons, which map onto distinct functional response types (Figure 3D, E), may influence the responses of other DH interneuron types and PSDC output neurons. We first assessed synaptic connectivity between primary afferents and PSDC output neurons, as well as the contributions of Rorβ and PVi inhibitory interneurons to feedforward and feedback inhibition using acute spinal cord slice electrophysiology (Figure 6A). Whole-cell voltage-clamp recordings were made from retrogradely labeled PSDC neurons, while optogenetically activating

**Figure 6.**
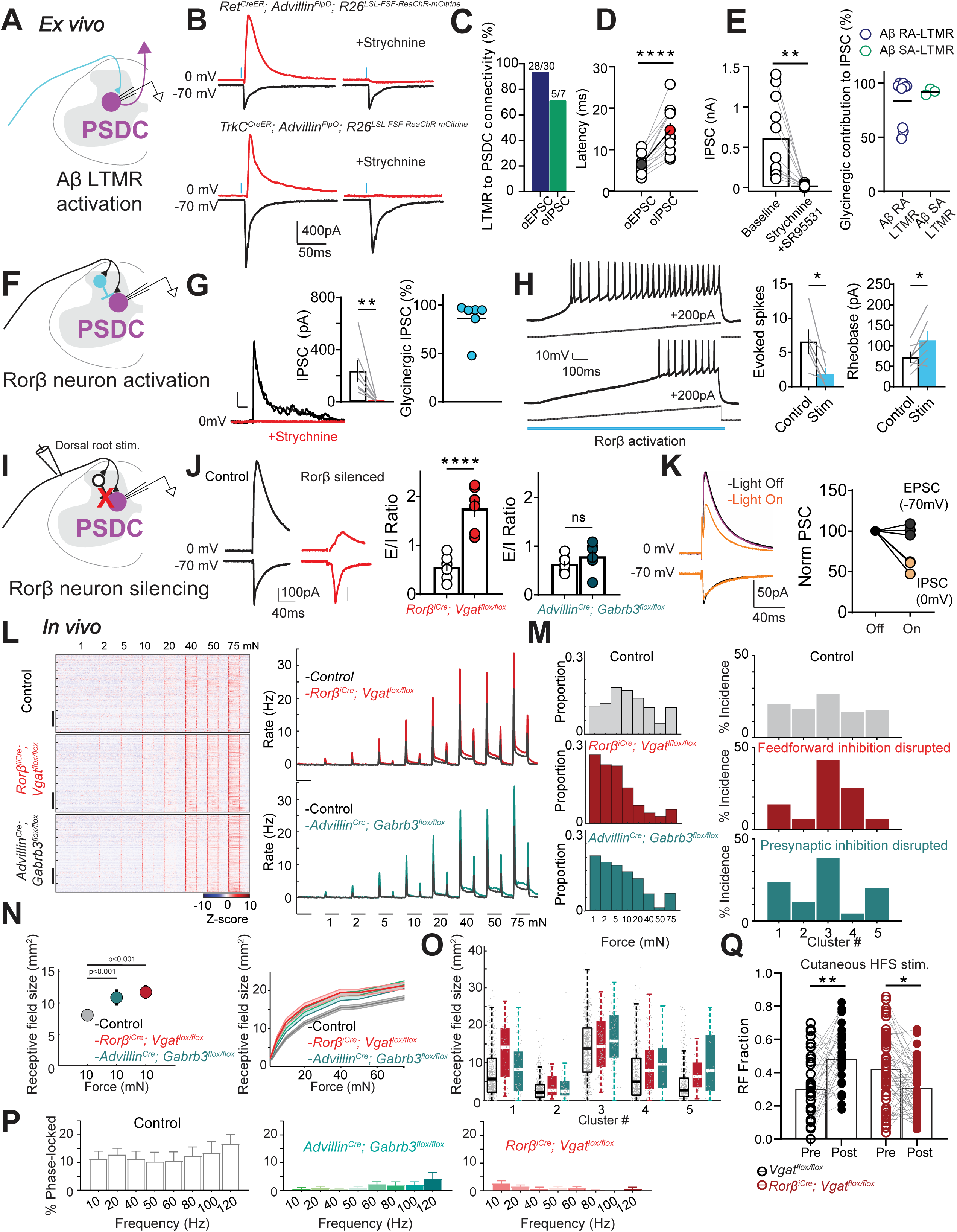
Sensory-evoked feed-forward and feedback inhibitory circuit motifs exert broad control over DH neuron responses. **A.** Recording configuration for optogenetic activation of LTMR synaptic terminals while recording from retrogradely labeled PSDC neurons in transverse lumbar spinal cord slices. **B.** Optical excitatory postsynaptic currents (oEPSCs) and polysynaptic inhibitory postsynaptic currents (oIPSCs) in retrogradely labeled PSDC neurons evoked by optogenetic activation of Aβ RA-LTMRs (using *Ret^CreER^; Advillin^FlpO^; R26^LSL-FSF-ReaChR::mCitrine^*mice) and Aβ SA-LTMR (using *TrkC^CreER^; Advillin^FlpO^; R26^LSL-FSF-ReaChR::mCitrine^* mice) synaptic terminals under baseline conditions and following application of the glycine receptor antagonist strychnine (2µM). **C.** Connectivity rates for Aβ RA-LTMRs and Aβ SA-LTMR oEPSCs to PSDC neurons. **D.** Onset latencies for opto-evoked Aβ RA-LTMR PSCs to PSDC neurons (n=14 PSDCs; N= 8 mice). Paired t test, P < 0.0001. Colored symbols represent the mean. Error bars: SEM. **E.** *Left*: Aβ RA-LTMR evoked oIPSC amplitudes before and after application of strychnine, and in some cases (n=3) of co-application of strychnine and the GABA_A_ receptor antagonist SR95531 (10uM) for the same PSDCs shown in D. Paired t two tailed test, P=0.002. Bars represent the mean. *Right*: Contribution of glycine and GABA to sensory evoked FFI. Percentage of IPSC component blocked by strychnine application in individual PSDCs (subsequent SR application blocked remaining strychnine insensitive component). Horizontal lines indicate mean values. P= 0.4818, Mann-Whitney *U* test. **F.** Schematic of experimental set-up. Optical activation of Rorβ neurons reduces excitability of PSDC output neurons. **G.** Light-evoked IPSCs from Rorβ neurons to PSDCs are blocked by bath-application of 2µM strychnine (n=7 PSDCs; N=5 mice). *Left*: representative glycinergic oIPSCs in PSDC neurons; Middle: oIPSC amplitudes before and after strychnine application. Paired t test, P < 0.05. *Right*: Contribution of glycine and GABA to Rorβ mediated IPSCs onto PSDC neurons. Strychnine insensitive IPSC component was blocked by application of SR95531. **H.** Current clamp recordings in PSDC neurons in response to 200pA current injection ramps paired with a 1s optical activation of Rorβ neurons. *Left*: Top trace shows evoked firing in PSDC neurons induced by the current injection; bottom trace shows the response in the same PSDC neuron when neighboring Rorβ neurons are coincidentally activated. *Middle*: Rorβ activation decreases the activity of PSDC neurons as indicated by a decrease in the total number of evoked spikes (quantified at 100pA current injection ± optical activation Rorβ neurons). Paired t test; n=6 neurons. *Right*: Rorβ neuron activation decreases the excitability of PSDC neurons as indicated by an increase in rheobase. Mann-Whitney U test; n=6 neurons. **I.** Schematic of experimental set-up and Rorβ silencing strategies used. **J.** Rorβ silencing in spinal cord slices from *Rorβ^iCre^; Cdx2-FlpO; R26^LSL-FSF-TeTox^* and *Rorβ^iCre^; Vgat^flox/flox^* mice reduced sensory-evoked FFI to PSDC neurons. *Left*: representative sensory- evoked EPSCs and IPSCs to PSDC neurons from transverse spinal cord slices with dorsal roots attached in littermate controls (black) and Rorβ silenced mice (red). *Middle*: E/I ratios are increased in Rorβ silenced mice (n=6; N=6 mice) compared to littermate controls (n=7; N=6 mice). Bars indicate mean ± SEM; P < 0.001; Mann-Whitney *U* test. *Right*: E/I ratios are similar between PSDCs from *Advillin^Cre^; Gabrb3^flox/flox^* mice (n=7 neurons; N=3 mice) and control littermates (n=5; N=4 mice). **K.** *Left:* Voltage-clamp recording from PSDCs isolates direct sensory-evoked EPSC (black) and polysynaptic IPSC (orange) at different holding potentials in spinal cord slices from *Rorβ^iCre^*; R26^LSL-eNpHR3.0-YFP^ mice. *Right:* Summary of normalized EPSC and IPSC amplitudes with (ON) and without (OFF) suppression of Rorβ neurons. N=3 mice; n=3 PSDCs; paired two-tailed t test. P>0.1 EPSC; P < 0.05 IPSC. **L.** *Right:* Z-scored firing rates for all dorsal horn units from control (top, n=486 units; N=14 mice), mice lacking Rorβ mediated FFI (FFI cKO: *Rorβ^iCre^; Vgat^flox/flox^* ; middle, n=474 units; N=12 mice), and from mice lacking PSI (PSI cKO: *Advillin^Cre^; Gabrb3^flox/flox^* ; bottom, n=507 units; N=12 mice). Units are sorted by depth. Scale bar: 100 units. *Left:* Baseline subtracted mean firing rate to a series of step indentations from 1-75mN across all control (grey), FFI cKO (red), and PSI cKO (teal) units. **M.** *Left*: Distribution of mechanical indentation thresholds in all recorded units from control (grey); FFI cKO (red), and PSI cKO mice (teal). *Right:* Incidence of DH functional response profiles in control (grey); FFI cKO (red) and PSI cKO mice (teal). Note that at a population level, the cluster distribution is uniform in control animals, but disrupted in FFI and PSI cKO animals. **N.** RF size measured at 10mN in all mechanically sensitive units from control (grey), FFI cKO (red) and PSI cKO (teal) mice. P < 0.001; standard deviation of bootstrapped RFs *Right*: RF areas across all forces tested in control (grey); FFI cKO (red) and PSI cKO (teal) animals. Mean ± SEM (shaded areas). **O.** RF areas measured at 10mN across all 5 functional response clusters in control (grey); FFI cKO (red), and PSI cKO animals. **P.** Phase-locked responses to mechanical vibrations are impaired in DH units from FFI cKO (red) and PSI cKO mice. **Q.** RFs size to 10mN mechanical step indentations increases following cutaneous electrical high frequency stimulation (HFS) in controls (black; N=2 mice), but not in mice lacking Rorβ mediated FFI (red; N=2 mice). WT: P < 0.01; Paired two-tailed t-test; FFI cKO: P < 0.05; paired two tailed t -test.

Aβ RA-LTMR or Aβ SA-LTMR terminals in spinal cord slices prepared from *Ret^CreER^; _Advillin_FlpO_; R26_LSL-FSF-ReaChR::mCitrine* _or *TrkC*_*CreER_; Advillin_FlpO_; R26_LSL-FSF-ReaChR::mCitrine* _mice,_ respectively. Photostimulation of Aβ RA-LTMR or Aβ SA-LTMR central terminals evoked large monosynaptic excitatory postsynaptic currents (EPSCs) in PSDC neurons (Figure 6A-C) that remained in the presence of TTX and 4AP (Figure S5A), consistent with their monosynaptic nature (Petreanu et al., 2009; Cruikshank et al., 2010). Moreover, selective activation of Aβ RA-LTMR or Aβ SA-LTMR synaptic terminals reliably elicited strychnine-sensitive polysynaptic inhibitory postsynaptic currents (IPSCs) in PSDC neurons through feedforward inhibition (FFI) in the local DH circuit (Figure 6B-E and Figure S5A). Optogenetic activation of Nav1.8^+^ sensory neuron terminals evoked smaller, polysynaptic EPSCs in PSDC neurons (Figure S5B).

We next assessed the contributions of Rorβ and PVi inhibitory interneurons to sensory neuron-evoked FFI in PSDC output neurons. Optogenetic stimulation of Rorβ interneurons (using both *Ror*β*^iCre^* and *Ror*β*^CreER^* driver lines) evoked strong monosynaptic IPSCs in PSDC neurons (Figure 6F, G), and, in current clamp experiments, directly suppressed the activity and excitability of neighboring PSDC neurons (Figure 6H; S5E). In contrast, optogenetic activation of PVi neurons, in slices from *PVi^Cre^;Lbx1^FlpO^; R26^LSL-FSF-ReaChR::mCitrine^* mice, did not evoke direct, monosynaptic IPSCs in PSDC neurons (Figure S5C). Consistent with these findings, immunohistochemistry in combination with retrograde viral labeling of PSDC neurons (RVdeltaG—GFP DCN injections in *Ror*β*^CreER^; R26^LSL-synaptophysin-tdTomato^* mice and gephyrin immunostaining) revealed that Rorβ interneurons in deep dorsal horn predominantly form inhibitory synaptic contacts onto the perisomatic compartment of PSDC neurons (Figure S5G). On the other hand, PVi neurons, which gate sensory input to the DH through inhibitory feedback (PSI), form specialized axo-axonic synaptic contacts onto sensory afferent terminals (Abraira et al., 2017; Boyle et al., 2019).

In related experiments, we used dorsal root electrical stimulation at intensities that selectively recruit Aβ fibers (Torsney and MacDermott, 2006; Methods) to measure somatosensory neuron-evoked EPSCs and polysynaptic IPSCs onto PSDC neurons in slices from mice in which distinct inhibitory interneuron subtypes were silenced. Rorβ neuron intersectional inactivation strategies, using either *Ror*β*^iCre^* in conjunction with *Cdx2-NSE-FlpO* for restricted FlpO expression to the spinal cord and the dual recombinase tetanus toxin line *RC:PFtox* or selective block of GABA/glycine release from Rorβ^+^ lineage neurons with *Ror*β*^iCre^; Vgat^flox/flox^* mice, were used to silence Rorβ interneurons. Recordings in spinal cord slices from these mice revealed that inhibiting Rorβ interneurons strongly attenuated Aβ-fiber evoked glycinergic FFI onto PSDC neurons (Figure 6I, J). In complementary experiments, acute optogenetic silencing of Rorβ neurons using *Ror*β*^iCre^; R26^LSL-Haporhodopsin-YFP^* mice selectively suppressed 50% of FFI onto PSDC neurons without affecting monosynaptic excitatory transmission from sensory afferents (Figure 6K). To assess the role of presynaptic inhibition (PSI) of primary afferent terminals to sensory-evoked EPSCs in PSDC neurons, which is mediated at least in part by axo-axonic synapses formed by PVi inhibitory interneurons onto primary afferent terminals (Abraira et al., 2017; Boyle et al., 2019), we used slices from mice that lack presynaptic GABA_A_ receptors in somatosensory neurons (*Advillin^Cre^; Gabrb3^flox/flox^* mice) and therefore lack low-threshold sensory-evoked GABA dependent PSI (Orefice et al., 2016; Zimmerman et al., 2019). In contrast to mice lacking FFI, genetic disruption of PSI did not alter sensory-evoked E/I ratios in PSDC neurons (Figure 6J). Together with previous findings (Boyle et al., 2019; Orefice et al., 2016; Zimmerman et al., 2019), these *in vitro* recordings indicate that Rorβ^+^ interneurons mediate the majority of sensory neuron-evoked FFI onto PSDC neurons, whereas PVi interneurons and presynaptic GABA_A_ receptors mediate low-threshold evoked GABA-dependent PSI of primary afferent terminals in the DH.

We next used these FFI and PSI genetic silencing strategies to assess the *in vivo* contributions of sensory-evoked inhibition, mediated by distinct interneuron populations, in shaping DH interneuron and PSDC projection neuron responses to mechanical stimuli. We first measured mechanical sensitivity, RFs, vibration tuning, and functional response cluster assignments in DH neurons using large scale MEA recordings, in the presence or absence of Rorβ-mediated FFI or GABA_A_ receptor-mediated PSI. Interestingly, disrupting FFI or PSI caused a dramatic increase in sensitivity to step indentations and sensory-evoked spiking responses in DH neurons (Figures 6L-M; S5J). Moreover, while inhibition of either Rorβ-mediated FFI or GABA_A_ receptor- mediated PSI shifted DH response profiles into a more sensitive range, disruption of FFI and PSI differentially altered the diversity of DH response cluster types (Figure 6M). Since the genetic inactivation strategies used are developmental and may lead to compensatory changes, we complemented these population level analyses with experiments in which Rorβ interneurons or PVi interneurons were acutely silenced using optogenetic strategies. Confirmatory *in vitro* slice recordings showed that acute Rorβ silencing (Figure S5D) indeed decreased sensory evoked FFI onto PSDC neurons (Figure 6K). As with the developmental deletions and population-level analyses, acute *in vivo* optogenetic silencing of Rorβ neurons during mechanical stimulation of glabrous skin increased indentation evoked spiking in neighboring DH units (Figure S5I).

Similarly, increased sensitivity to step indentations was observed in neighboring units following acute optogenetic silencing of PVi neurons (Figure S5H).

We also assessed DH neuron RF properties in the absence of Rorβ interneuron mediated FFI and GABA_A_ receptor-mediated PSI. Inactivation of FFI and PSI similarly increased RFs across all recorded DH neurons and different cluster types (Figures 6N, O). Differences between the FFI and PSI groups did emerge, however, upon analyzing spatiotemporal features of DH neuron RFs. While RFs measured at the ON response are significantly larger in both the FFI and PSI mutants (Figure 6N), RFs measured during the sustained and OFF portions of indentation steps in the FFI mutants, but not the PSI mutants, differed from controls (Figure S5K). Moreover, DH neuron phase-locking to vibratory stimuli was nearly abolished in the absence of FFI or PSI (Figure 6P), while vibration frequency dependence of force sensitivity across DH neurons and all cluster types was minimally affected in both mutants.

Finally, the potential contribution of one form of sensory-evoked inhibition, Rorβ interneuron mediated FFI, to high frequency stimulation evoked plasticity of DH responses was tested.

Remarkably, while high frequency electrical stimulation of the tibial, sural, and saphenous nerves evoked rapid and marked increases in both the sustained responses and RF properties across a large cohort of DH neurons, such stimulus-evoked increases in DH neuron response properties were absent in FFI mutant mice (Figures 6Q and S5L-N). Related to this, in spinal cord slice recordings with dorsal roots attached, high frequency stimulation triggered robust long-term potentiation (LTP) of Aβ sensory neuron synapses onto PSDC output neurons, and this form of plasticity was absent in slices from mice lacking Rorβ interneuron mediated glycinergic FFI (Figure S5F).

Together, these findings indicate that FFI and PSI, and thus the interneurons within clusters 1 and 3 that mediate these forms of synaptic inhibition, shape the mechanical responses of most and possibly all DH neurons, across all functional cluster types. This supports a model in which DH interneuron types are functionally inter-dependent: interneurons of one functional type shape the mechanical responses and RFs of other DH interneuron types. Moreover, DH neuron response properties and RFs across the functional cluster types flexibly adjust to high frequency volleys of sensory inputs, and at least one DH inhibitory motif, glycinergic FFI mediated by the Rorβ interneurons of cluster 1, is required for this widespread sensory-evoked plasticity in the DH.

### PSDC output signals reflect convergent mechanosensory inputs and are shaped by distinct modes of DH synaptic inhibition

We next tested the hypothesis that DH output signals conveyed by PSDC neurons reflect the extensive convergence of mechanoreceptor inputs and the inter-dependency of DH circuit components. Thus, we selectively recorded from PSDC neurons using an optotagging strategy and recording configuration that enables *in vivo* measurements of tactile-evoked responses in PSDC neurons in conjunction with sensory neuron and DH neuron functional manipulations (Figure 7A; Figure S6A-F; see Methods). Overall, tagged PSDC neurons exhibited a surprisingly high diversity of response properties and sensitivities to mechanical step indentations (Figure 7B, D). Some PSDCs exhibited very low sensitivity (1 mN force thresholds), others had higher force thresholds, and some displayed prominent sustained responses and encoded forces well into the HTMR range (Figure 7B, Control). PSDC neurons also displayed variably shaped and typically large cutaneous RFs (Figure S6I). Thus, unlike the genetically labeled DH interneuron types, which exhibited stereotyped responses to simple tactile stimuli, PSDC neurons exhibit a strikingly large range of sensitivities, firing patterns, and RF areas (Figure 7B, D, E). We also examined PSDC neuron responses to sinusoidal vibrations of varying amplitudes and frequencies and found that phase-locking to mechanical vibrations exceeding 20Hz was absent in these projection neurons (Figure S6J), although PSDCs did display vibration frequency-dependent sensitivity suggesting convergent inputs from Aβ RA1-LTMR (Meissner) and Aβ RA2-LTMR (Pacinian) signals (Figure S6J). To directly address the extent of Aβ LTMR subtype convergence onto PSDC neurons, we employed a complementary tagging strategy and used electrical antidromic stimulation of PSDC dorsal column axons at cervical levels C1-C3 in mice expressing ReaChR in either Aβ RA-LTMRs or Aβ SA-LTMRs (Figure S6G, H). Selective activation of Aβ RA-LTMRs or Aβ SA-LTMRs evoked action potentials in most mechanically sensitive PSDC neurons tested demonstrating convergent input from both of these Aβ LTMR subtypes (Figure S6H).

**Figure 7.**
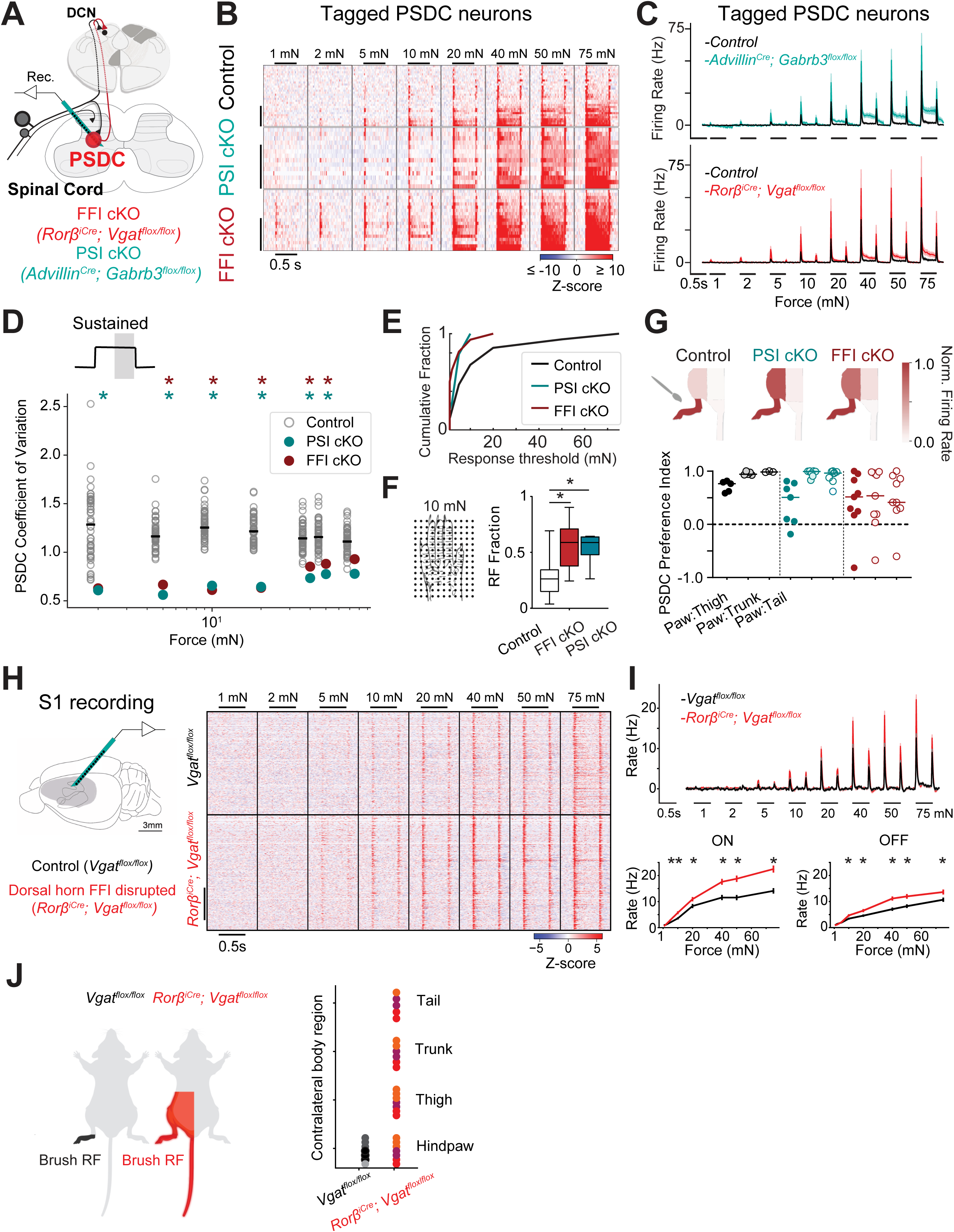
DH synaptic inhibition shapes response properties and receptive fields of PSDC output neurons and neurons in S1. **A.** Schematic of experimental set-up. **B.** Heatmaps of z-scored firing rates for tagged PSDC units from control (n=34; N=18 mice); FFI cKO (*Rorβ^iCre^; Vgat^flox/flox^* ; n=17; N=6 mice) and PSI cKO (*Advillin^Cre^; Gabrb3^flox/flox^* ; n=14; N=5 mice) animals. Units are sorted by the magnitude of their sustained response. **C.** Baseline subtracted grand mean firing rate (±SEM) in tagged PSDC neurons across all three genotypes (black traces, pooled PSDC neurons in control mice, red traces, PSDC neurons in FFI cKO mice; teal traces, PSDC neurons in PSI cKO mice). Error bars are SEM and center values are grouped means. **D.** C.V. of sustained response magnitudes at indentation force steps between 2 to 75mN, measured from PSDC neurons in FFI cKO (red markers; n=17); PSI cKO (teal markers; n=14) and 50 representative sets of 14 randomly sampled PSDC neurons from control mice (grey markers). Asterisks denote parameters for which the value in FFI cKO and PSI cKO PSDC neurons was lower than that for WT PSDC neurons in <0.5% of 10 000 subsamples (i.e., p < 0.05). **E.** Cumulative distributions of mechanical indentation thresholds in control PSDC neurons compared to PSDCs from FFI cKO or PSI cKO mice. WT vs. FFI PSDC neurons: P < 0.005; WT vs. PSI PSDC neurons P < 0.05, Mann-Whitney *U* test. **F.** RF fraction of skin area mapped with10mN force steps in control PSDC neurons vs. PSDC neurons from FFI cKO and PSI cKO animals. P < 0.01, Mann-Whitney *U* test. **G.** Brush-evoked RFs in PSDC neurons from control (grey), FFI cKO (red) and PSI cKO (teal) animals. Spikes were evoked by stroking equivalent skin areas of the hindpaw, thigh, trunk, and proximal tail. Top: Representative responses to touch of the body in PSDC units from control, FFI cKO and PSI cKO mice. Firing rates normalized to the maximum brush-evoked response are depicted on the mouse schematic for an example PSDC neuron across all three genotypes. Bottom: PSDC body region preference index for a subset of PSDC units (computed as (FR_paw_ – FR_thigh_)/ (FR_paw_ + FR_thigh_); (FR_paw_ – FR_trunk_)/ (FR_paw_ + FR_trunk_); (FR_paw_ – FR_tail_)/ (FR_paw_ + FR_tail_)). For (FR_paw_ – FR_trunk_)/ (FR_paw_ + FR_trunk_) shows larger RFs in PSDCs from animals lacking FFI (n=9 PSDCs from 5 mice) or PSI (n=7 PSDCs from 4 mice) as compared to PSDCs from control animals (n=5 PSDCs). P < 0.05, Kruskal-Wallis H test. **H.** Left: Recording configuration for S1 *in vivo* electrophysiology in awake, head-fixed mice. An acute 32 channel silicon probe was implanted in the hindpaw region of S1. Right: Heatmaps of z-scored firing rates in hindpaw S1 from control *Vgat^flox/flox^* (top; n=335 units from 7 recording in 3 mice) and *RorβiCre; Vgat^flox/flox^* mice (bottom; n=373 units from 7 recordings in 3 mice). Regular spiking and fast spiking units are combined in this analysis and sorted by depth. **I.** Top: grand mean (±SEM) baseline subtracted firing rates to step indentations across all control (grey) and FFI cKO (red) units. Bottom: tuning curves i.e. the mean (±SEM) baseline subtracted firing rate at the step onset (left) and offset (right) for each force in units from wild-type (grey) and FFI cKO (red) animals. **J.** Schematic of brush-evoked RF in hindpaw S1 units from control and FFI CKO animals. Right: Brush-evoked responses across multiple body regions in recordings from control (dark grey hues) and FFI cKO (red hues) mice. Multiple recordings from the same animal are represented with markers of the same color (within a genotype).

We next evaluated the contributions of Rorβ interneuron mediated FFI and GABA_A_ receptor mediated PSI, respectively, in shaping the response properties and RFs of PSDC neurons.

Disrupting either FFI or PSI decreased mechanical thresholds and increased indentation-evoked spiking in PSDC neurons (Figures 7B, C, E). Spiking responses at the ON and OFF portions of step indentations were elevated across the entire force range, demonstrating that both FFI and PSI gate Aβ RA-LTMR and Aβ SA-LTMR drive onto PSDC output neurons (Figures 7B, C, S7A). Responsivity during the sustained phase of step indentations was strongly elevated at indentation forces that recruit HTMR input, and to a greater extent in mice lacking PSI than mice lacking FFI (Figures 7B, C and S7A). This is consistent with the finding that PVi interneurons exhibit more robust sustained responses to indentation compared to Rorβ interneurons (Figures 3H-J). Furthermore, a prominent after-discharge following step indentations was observed in mice lacking PSI, but not FFI, indicating the PSI uniquely contributes to PSDC response kinetics and temporal filtering of mechanosensory inputs (Figure 7B, C). Both forms of synaptic inhibition also shape PSDC RFs on glabrous hindpaw: RFs mapped with 500 ms indentation steps were substantially larger in the absence of either FFI or PSI across all force intensities tested (Figures 7F, S7C). PSDC RFs outside of the hindpaw region were also mapped using brush stimuli (Figure 7G). Strikingly, while PSDC neurons in control mice showed a strong preference for hindpaw stimulation compared to thigh stimulation, mutants lacking PSI exhibited comparably strong responses to stroking the hindpaw and thigh (Figure 7G). Similarly, PSDCs in mice lacking FFI responded to stroking across the hindpaw and thigh, and in some cases back hairy skin and proximal tail stroke, revealing large, multiregional receptive fields (Figure 7G).

Finally, while PSDC neurons in control mice exhibited highly diverse response properties and sensitivities, as observed in the heat maps of individual PSDC responses and quantified using a coefficient of variation analysis of the entire population (Figure 7D, S7B), PSDCs in mice lacking FFI or PSI had considerably less variation in their mechanical response thresholds, response profiles, and RFs (Figure 7D, E). Thus, DH interneurons that mediate distinct modes of DH inhibition and control the mechanical responses of all other DH interneuron types are also crucial in shaping the wide diversity of responses observed across the PSDC neuron population.

### DH outputs dictate tactile responses at supraspinal regions

Because PSDC neurons convey tactile information from the DH to the DCN, and as a group exhibited heightened sensitivity and larger RFs in both the FFI and PSI mutant mice, we hypothesized that in the absence of FFI or PSI in the DH: *(1)* downstream neurons will exhibit increased sensitivity and response amplitudes to skin indentation; *(2)* touch-evoked RFs will be expanded; and *(3)* behavioral responses to tactile stimuli will be altered. We chose the Rorβ interneuron mediated FFI disruption model to test these predictions because the genetic manipulations used to disrupt FFI are restricted to the spinal cord (Figure S7F), whereas the PSI disruption model is likely to eliminate GABA_A_ receptor mediated PSI of primary afferent terminals in the DCN thereby disrupting the direct dorsal column pathway. Therefore, we next recorded touch-evoked responses in S1 (Emanuel et al., 2021) of awake *Rorβ^iCre^; Vgat^flox/flox^* mice and *Vgat^flox/flox^* littermate controls. We found that selective disruption of Rorβ neuron mediated glycinergic FFI in the spinal cord increased spike rates at both the onset and offset of step indentations, without altering S1 transient response profiles (Figures 7H, I, S7D,E). We also found a striking alteration of RF properties of individual S1 units in the absence of FFI in the DH. While units from hindlimb S1 responded most robustly to hindlimb skin stroking in control mice, individual S1 units recorded in *Rorβ^iCre^; Vgat^flox/flox^* mice responded comparably to stroke stimuli applied to many regions spanning the entire contralateral lower body (Figure 7J).

Consistent with these alterations in the physiological properties and RFs recorded in S1, behavioral measurements showed that disruption of Rorβ neuron mediated FFI in the DH caused tactile over-reactivity (Figure S7H), deficits in texture discrimination (Figure S7I), and an alteration in a sunflower seed handling behavioral assay that assesses the integrity of sensory-motor exchange and dexterous use of the forepaws (Figure S7J). Together, these findings indicate that mechanoreceptor signal processing in the DH shapes touch-evoked responses and topographic body representation at supraspinal levels as well as somatosensory behaviors.

## Discussion

Here we combined large-scale *in vivo* electrophysiology with a range of functional manipulations to assess spinal cord DH interneuron and output neuron response type diversity and function with the goal of extracting principles of mechanosensory information processing at the earliest stage of the somatosensory system hierarchy. Our findings support a model in which an array of functionally distinct DH interneuron types receive extensively convergent LTMR and HTMR inputs and form a highly inter-connected network architecture that functions to flexibly shape a diversity of response properties of PSDC output neurons. We also provide evidence that mechanosensory signal transformations within the intricate DH network dictate how mechanical forces acting on the skin are represented in the brain.

### Functional classification of dorsal horn neurons encoding touch

DH neurons have been described based on a range of properties, including expression of immunohistological and genetic markers, morphological and anatomical features, and intrinsic physiological signatures, however *in vivo* response and RF properties of DH neuron types are poorly understood or not known. Our large-scale electrophysiological analysis and unsupervised clustering complements prior analyses and revealed six functional types of DH neurons based on their responses to innocuous touch stimuli acting on mouse glabrous skin. These functional groups exhibit distinct sensitivities, tonic and mechanically evoked firing patterns, and RF sizes and shapes. Most encode mechanical force over a wide range of intensities and are also sensitive over a broad range of vibration frequencies. A sampling of genetically labeled interneuron types indicates that previously identified, genetically defined subtypes map mainly onto one or a few of the distinct functional classes observed here. Future studies combining similar large scale *in vivo* recordings and stimuli that span other somatosensory modalities, including thermal and chemical space, will further define the functional properties of the DH neuron types described here.

### Extensive LTMR subtype and HTMR convergence in the dorsal horn

#### Aβ RA-LTMRs and Aβ SA-LTMRs converge onto most or all functionally defined DH neuron types

Several lines of evidence point to Aβ LTMR subtype convergence onto the majority of DH neurons across the full range of DH neuron functional types. *First*, the majority of mechanically sensitive DH units across all functional response types have large and spatially complex cutaneous RFs when probed with 10mN force steps, which preferentially activates LTMRs, and these DH responses are completely absent in mice lacking both Meissner corpuscles and Merkel cells. Since the RFs of Aβ RA-LTMRs and Aβ SA-LTMRs themselves are considerably smaller than even the smallest DH neuron RFs, and the step indentations used here do not activate Pacinian LTMRs (Emanuel et al., 2021), DH RFs measured at low indentation forces must reflect homotypic and/or heterotypic convergence of signals from many individual Aβ RA-LTMRs and Aβ SA-LTMRs. *Second*, genetic disruption of both Aβ RA-LTMRs and Aβ SA-LTMR signals resulted in loss of responsivity to 10mN indentations across all mechanically sensitive DH units, indicating broad contribution of these Aβ LTMR types to virtually all DH responses in the low force range. Moreover, disruption of Aβ RA-LTMR signals alone led to loss of responsivity to low force indentations, and although OFF responses, characteristic of Aβ RA-LTMR input, were absent in the low force range, OFF responses at higher indentation forces were observed. This finding suggests that DH OFF responses are partially inherited from the periphery (i.e., driven by the Aβ RA-LTMR OFF response) and also generated *de novo*; a proposed mechanism for this is rebound from inhibition upon termination of slowly adapting mechanoreceptor types, in the high force range. *Third*, selective optogenetic activation of Aβ RA-LTMR or Aβ SA-LTMR terminals in glabrous skin to evoke one action potential in a few Aβ LTMRs was sufficient to evoke spiking across the majority of DH neurons representing all functional response types, as well as in PSDC projection neurons. This observation is particularly striking when considering that activation of Aβ SA-LTMRs evoked spiking even in DH neurons that responded transiently to mechanical step indentations, while activation of Aβ RA-LTMRs evoked spiking in DH units with sustained mechanical response profiles. Collectively, these findings demonstrate extensive Aβ RA-LTMR and Aβ SA-LTMR signal convergence and non-linear transformations onto most if not all mechanically sensitive DH neurons, giving rise to distinct DH interneuron and output neuron firing patterns.

#### Vibration encoding in the DH suggests extensive Aβ RA1-LTMR and Aβ RA2-LTMR convergence

As in the somatosensory periphery, DCN and cortex, a select subset of DH neurons exhibited phase-locked spiking in response to mechanical vibrations (sinusoids) delivered to hindpaw glabrous skin. PKCγ^+^ interneurons are unique among the DH populations sampled here because a large fraction of this population exhibited phase-locking to mechanical vibrations up to 120Hz. Interestingly, while relatively few DH neurons are phase-locked above 20 Hz, the vast majority of DH neurons across all functional response types exhibited broad vibration frequency dependent tuning (i.e., frequency dependent sensitivity to skin indentation). Together with the observation that DH units from *TrkB^cKO^* mice lack vibration tuning in the Meissner range, but not in the Pacinian range, these findings suggest that most DH neurons receive input, either directly or indirectly, from both Meissner corpuscle Aβ RA1-LTMRs and Pacinian corpuscle Aβ RA2- LTMRs, which exhibit distinct frequency selectivity (20-100 Hz and 60-500 Hz, respectively).

#### Convergence of LTMR and HTMR inputs across the DH broadens the dynamic range of DH outputs

Our gain-of-function and loss-of-function sensory neuron manipulation experiments also indicate that most DH neurons across all DH neuron functional response types receive inputs from HTMRs, either directly or indirectly. Indeed, we found that responses of most DH units do not saturate or plateau in response to indentations using force amplitudes at which most LTMRs saturate. Partial disruption of HTMR signals supports this notion, as do optogenetic sufficiency experiments demonstrating that A-fiber HTMR evoked responses are broadly distributed across the six principal functional response types. It is also noteworthy that optogenetic stimulation latency measurements indicated that most DH neurons receive convergent A-fiber and C-fiber inputs, in a topographically aligned manner. The surprisingly high degree of LTMR and HTMR convergence is also reflected at the level of PSDC output neurons, as the targeted *in vitro* and *in vivo* PSDC electrophysiological recordings demonstrate. Thus, one consequence of extensive LTMR and HTMR signal convergence in the DH is that the indirect dorsal column pathway encodes intensity of mechanical forces acting on the skin over a very wide dynamic range, as compared to the direct dorsal column pathway.

### Feedforward and feedback inhibitory motifs broadly shape tactile-evoked responses in the DH, revealing a highly interconnected network architecture

How are LTMR and HTMR inputs to the DH transformed into output signals conveyed to other CNS regions, including the DCN? At one end of the range of possibilities, the DH could be comprised of multiple, discrete channels, with individual or groups of interneuron subtypes shaping the response properties and RFs of select output populations. At the other end of this range, the DH could be an interconnected network of inter-dependent circuit elements, with each principal interneuron type uniquely contributing to the tactile responses of all other interneuron and output populations. Our findings favor the latter. DH Rorβ inhibitory interneurons, which fall mostly into functional class 1, mediate sensory-evoked glycinergic FFI in the LTMR-RZ. On the other hand, PVi inhibitory interneurons, which fall mostly into functional class 3, form a majority of axo-axonic contacts with central terminals of primary afferents in lamina II_i_, III and IV and participate in GABA_A_ receptor-dependent PSI (Abraira et al., 2017; Boyle et al., 2019; Hughes et al., 2012). Importantly, selective genetic perturbations and loss-of-function experiments revealed that both Rorβ-mediated FFI and GABA_A_ receptor-mediated PSI govern the sensitivity of neurons across all DH functional response profiles and PSDC output neurons. These findings are consistent with prior studies implicating FFI in the somatosensory cortex in controlling the dynamic range of cortical network responses (Pouille et al., 2009) and roles for PSI in general gain control in the retina (Nagy et al., 2021) and odor-evoked responses in olfactory circuits (Nunes and Kuner, 2015; Olsen and Wilson, 2008; Root et al., 2008).

Interestingly, PSI, but not FFI, governs temporal filtering of PSDC responses to step indentation, through preventing after-discharge spiking following the offset of step indentations. Loss of FFI or PSI also resulted in remarkably large, aberrant RFs across all functional classes of DH interneurons as well as PSDC output neurons. Together, these findings point to a highly connected, inter-dependent network model of the DH in which interneurons of distinct functional classes, and the circuit motifs they engage, directly or indirectly shape responses of all DH interneuron types of the network as well as PSDC output neurons.

### The DH and its role in mechanotransmission

To what extent does mechanosensory information processing in the DH dictate how touch is represented in the brain? Models to explain the functional organization of the discriminative touch pathway and the brain’s representation of innocuous mechanical stimuli acting on the skin have mainly focused on the direct dorsal column pathway, comprised of Aβ LTMR axonal projections extending via the dorsal column to the DCN. However, compared to the DCN, we estimate that the spinal cord DH receives between 10 and 100 times more primary mechanosensory neuron synapses, including the majority of Aβ LTMR synapses and possibly all Aδ fiber and C fiber mechanosensory neuron synapses. Moreover, the DH contains PSDC neurons whose axons form the indirect dorsal column pathway and ascend in parallel with direct dorsal column pathway axons to the DCN. What is the functional significance of the highly interconnected DH mechanosensory circuitry, with its broadly convergent LTMR and HTMR inputs and PSDC/indirect dorsal column pathway outputs? One clue may come from the response properties of PSDC neurons. We found that PSDC neurons, as a population, exhibit highly heterogeneous responses to tactile stimuli. Indeed, mechanical force thresholds, OFF responses, and magnitudes of sustained responses to step indentations as well as RFs are highly variable across the PSDC population. This functional diversity of PSDC output neurons is consistent with classic PSDC recordings from the cat and is reminiscent of the broad, heterogeneous tuning and RF properties of output neurons of other early sensory processing areas, including retinal ganglion cells (Baden et al., 2016; Emanuel et al., 2017; Milner and Do, 2017; Sanes and Masland, 2015). A second clue comes from the striking observation that this broad range of PSDC neuron response properties collapses into more homogeneous responses across the PSDC population in mice lacking FFI or PSI. Indeed, in the absence of FFI or PSI, most PSDC neurons responded to the lowest indentation forces tested, exhibited pronounced sustained firing patterns, particularly in the HTMR range, and displayed extremely large RFs. As such, proper DH network function is essential for setting the wide range of sensitivities, response properties, and RFs across the PSDC output neuron population. It is interesting to note that both *Rorβ* and *Gabrb3* are ASD associated genes (Iossifov et al., 2014; Satterstrom et al., 2020), raising the possibility that alterations in DH output diversity and flexibility may contribute to aberrant tactile behaviors observed in some genetic mouse models of ASD (Orefice et al., 2019; Orefice et al., 2016). A third clue comes from our observation that bursts of high frequency afferent stimulation evoked changes in the response properties and RFs of neurons across the DH, which is consistent with the observation from A.G. Brown and colleagues (Brown et al., 1983) of activity-dependent changes in PSDC RF properties. Also noteworthy are our observations that afferent stimulation evoked changes in DH neuron response properties and activity dependent potentiation of Aβ LTMR-PSDC neuron synapses are both eliminated in mice lacking FFI. That the DH is modulated by top-down control via corticospinal inputs, and corticospinal inputs to the DH distribute broadly across genetically defined DH interneuron types, is also of interest when considering models of DH network function. Collectively, these observations lead us to propose a model in which the highly interconnected DH circuitry, with its broadly convergent LTMR and HTMR inputs, functions to enable a wide, flexible range of DH interneuron and PSDC output neuron sensitivities, firing patterns and RF properties, modifiable by sensory experience and internal state (e.g., movement, attention). Future experiments will test how mechanosensory processing in the DH and its variable, modifiable PSDC output signals combine with unmodified LTMR signals carried by the direct dorsal column pathway to shape the brain’s representations of stimulus intensity, texture, direction and speed of stimulus movement, object orientation, softness, and compliance, among other features of the physical world.

## Acknowledgements

We thank Ofer Mazor and Pavel Gorelik from the HMS Research Instrumentation Core Facility technical assistance, Danielle T. Morency, Terri Javaluyas, Kaitlyn Clausel, Sarah Tsan, and Matthew Yee for assistance with mouse husbandry and immunohistochemistry, Michelle M. DeLisle for assistance with behavioral experiments, and all members of the Ginty lab for helpful comments on the manuscript. This work was supported by postdoctoral fellowships from The Harvard Mahoney Neuroscience Institute Fund (AMC) and The Ellen R. and Melvin J. Gordon Center for the Cure and Treatment of Paralysis (AMC), predoctoral fellowships NSF GRFP DG1745303 (GR) and Stuart H.Q. & Victoria Quan Fellowship (S-YT), NIH grants DP1 MH125776 (CDH), R01 NS089521 (CDH), NS119739 (AJE), and NS097344 (DDG), The Hock E. Tan and Lisa Yang Center for Autism Research at Harvard University (DDG), and the Edward R. and Anne G. Lefler Center for Neurodegenerative Disorders (DDG). DDG is an investigator of the Howard Hughes Medical Institute. This article is subject to HHMI’s Open Access to Publications policy. HHMI lab heads have previously granted a nonexclusive CC BY 4.0 license to the public and a sublicensable license to HHMI in their research articles. Pursuant to those licenses, the author-accepted manuscript of this article can be made freely available under a CC BY 4.0 license immediately upon publication.

## Author contributions

AMC, GR, and DDG conceived the study. AMC and GR performed all of the *in vivo* spinal cord electrophysiological experiments. AMC, GR and AJE performed *in vivo* DRG recordings. AJE performed S1 *in vivo* electrophysiology experiments. AMC performed *in vitro* slice electrophysiology. CCM and DZ performed anatomical experiments with assistance from AMC. AMC and GR analyzed data with assistance from S-YT, AE, and CDH. AMC, GR and DDG wrote the paper with input from all authors.

## Supplemental figure legends

**Figure S1.**
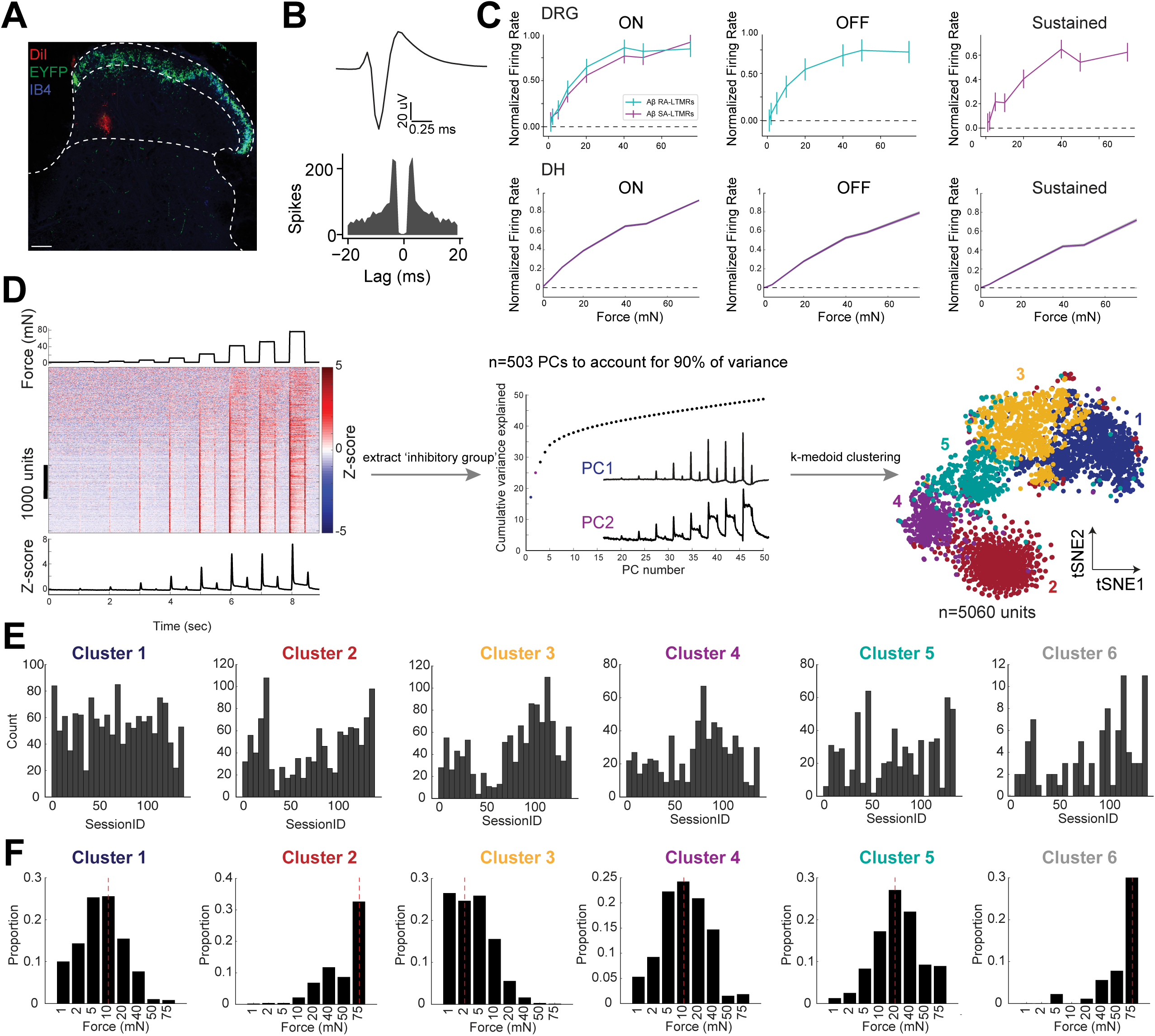
Functional characterization of mechanosensory responses in DH neurons. **A.** Example DiI-coated MEA in L4 spinal DH from a mouse expressing ChR2-YFP in Mrgdprd^+^ polymodal nociceptors (*Mrgdprd^CreER^; R26^LSL-ChR2-EYFP^*; green, EYFP; blue, IB_4_ binding protein; red, DiI coating probe tract). Scale bar, 100μm. **B.** Average waveform and corresponding autocorrellogram for example DH unit. **C.** Top: Average maximum-normalized responses (±SEM) for Aβ SA-LTMRs (n = 11 neurons) and Aβ RA-LTMRs (n = 13 neurons) at the ON, OFF, and sustained portion of step indentations. Note saturation of all response phases around 40 mN. Bottom: Average maximum-normalized firing rate for all mechanically sensitive DH neurons. Unlike DRG neurons, DH neurons do not plateau at these force steps. Shaded regions indicate SEM. **D.** Unsupervised clustering pipeline for identifying DH functional response profiles. Left: Heatmap of individual DH units responding to indentation steps. Grand average z-scored PSTH is depicted below. First, units with decreased firing rates to indentations steps, ‘inhibitory group’, were separated from the overall dataset. For dimensionality reduction, PCA analysis was then used to extract features of indentation-evoked DH responses from the remaining dataset. Inset shows PC1 and PC2 accounting for 25% of variance in the data, and resembling responses with transient or sustained components. Finally, we used the number of PCs accounting for 90% of the variance (n=503) for k-mediods clustering. This approach yields 6 principal functional clusters, (right) 5 clusters shown in TSNE space and the 6^th^ is comprised of the ‘inhibitory group’. See Methods for details. **E.** Occurrence of units sorted into the 6 principal clusters across experiments. **F.** Proportion of unit response thresholds within each principal cluster (median is denoted as dashed red line).

**Figure S2.**
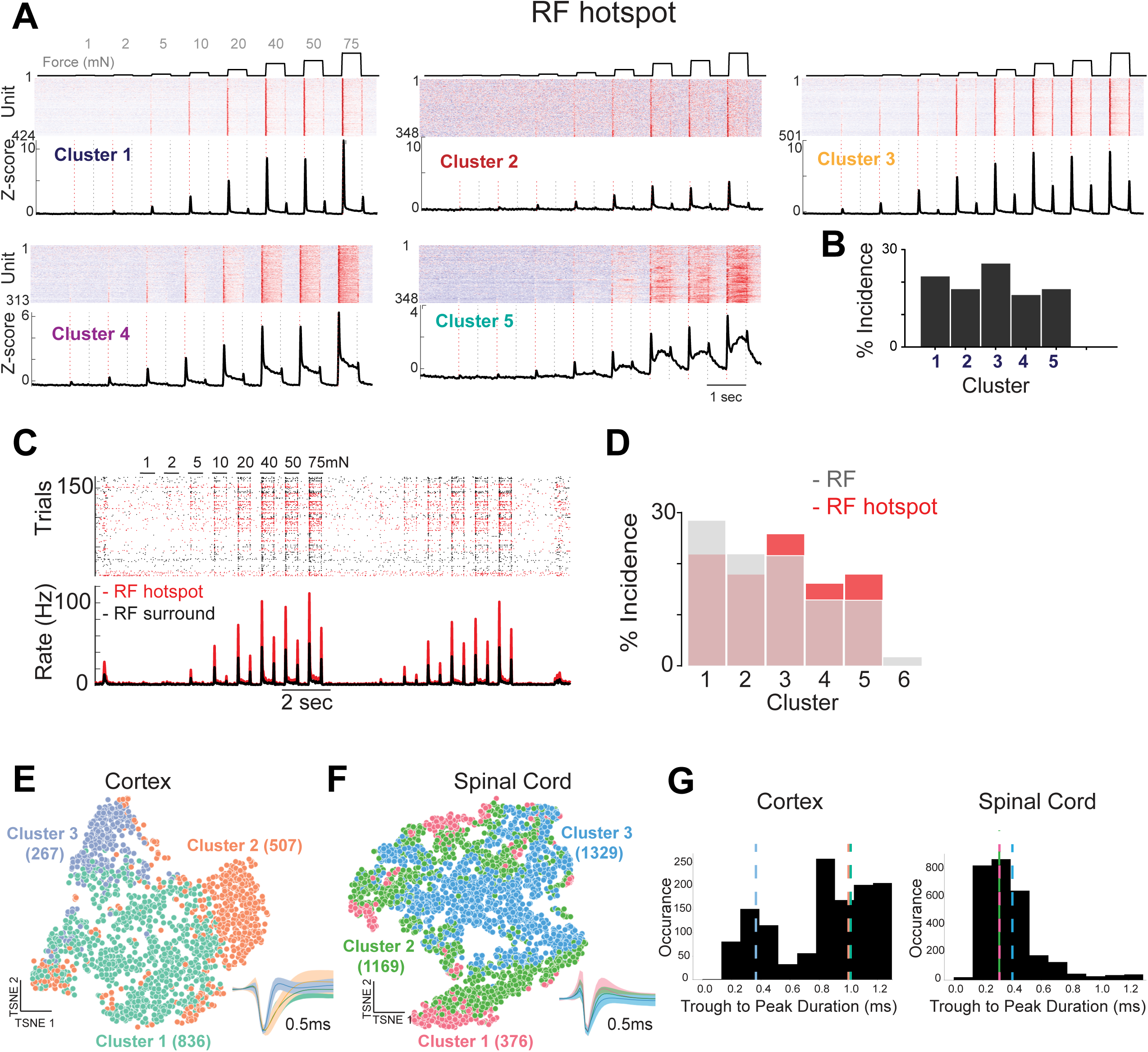
Functional response profiles at RF hotspot and extracellular waveform features. **A.** Functional clustering was performed as described (see Methods; Figure S1 D) at the RF hotspot. For each cluster, a heatmap of all included DH units responding to indentation is shown above the average z-scored PSTH. Heatmaps are sorted by threshold. **B.** Percentage of DH units used in RF hotspot clustering per functional cluster. **C.** Spike raster plot of DH unit linear RF organization with trials in the RF hotspot colored in red (top). Mean PSTHs for RF hotspot (red) and RF surround (black) shown below. Note the increase in firing rate from RF surround to RF hotspot. **D.** Percentage of units in the total dataset (n=5060), using all RF information, per functional cluster (grey) compared to the percentage of units used in the RF hotspot clustering (red). **E.** t-SNE visualization of somatosensory cortex extracellular waveforms clustered using k-means clustering (n= 1610 units; see Methods). As previously reported, three primary waveform subtypes emerged: two regular-spiking (green, cluster 1; orange, cluster 2) and one fast-spiking (purple, cluster 3). Normalized mean waveforms in different clusters are shown ±SD. **F.** DH extracellular waveforms (n = 2874 units) clustered as in (A). Unlike cortex, DH waveforms are not clearly separated into three groups (pink, cluster 1; green, cluster 2; blue, cluster 3). Normalized mean waveforms for units in different clusters are shown ±SD. **G.** Trough-to-peak durations of cortical (left) and DH (right) units. Cortical waveforms have a bimodal distribution, highlighting the duration difference between fast-spiking (dashed purple line, cluster 3; median: 0.35ms) and regular-spiking (dashed green line, cluster 1; median: 1.05ms; dashed orange line, cluster 2; median: 1.00ms) units. Spinal cord waveforms have a unimodal distribution (dashed pink line, cluster 1; median: 0.30ms; dashed green line, cluster 2; median: 0.30ms; dashed blue line, cluster 3; median: 0.40ms).

**Figure S3.**
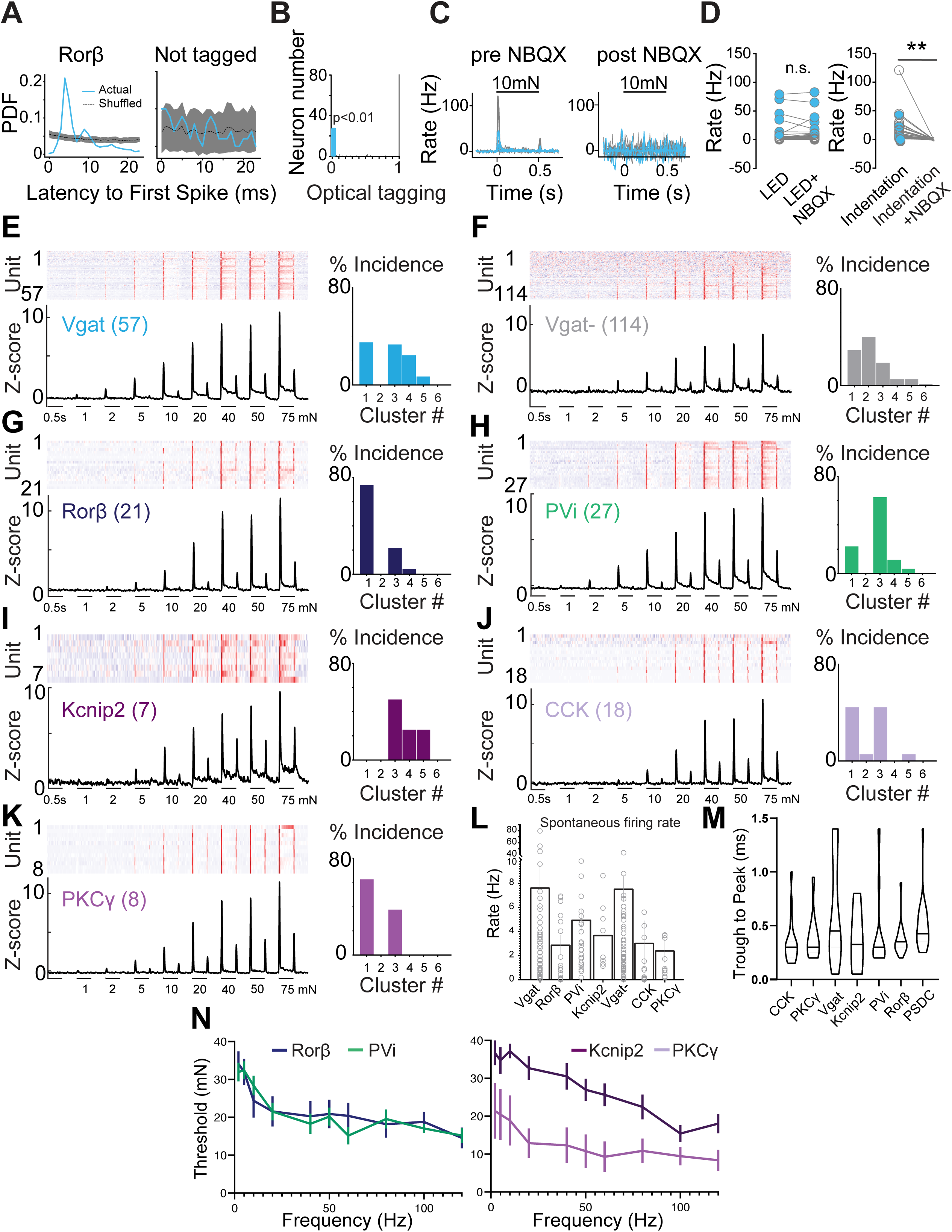
Functional characterization of distinct DH interneuron types. **A.** Light-evoked first spike latency probability distribution from the optotagged Rorβ^+^ unit in B (left) and non-tagged neighboring unit (right). **B.** Histogram of optical test (SALT) P values for optically tagged Rorβ^+^ units and simultaneously recorded non-tagged units (n=21; 5 mice; p < 0.01, blue for tagged units; p=1; grey for non-tagged units). **C.** Raster plot and PSTH of the same Rorβ^+^ unit in the absence or presence of NBQX (5 mM) applied to the surface of the cord to block excitatory synaptic transmission. **D.** Summary light-evoked firing before and after NBQX application. Optotagged units (blue) had similar optically evoked firing rates in the two conditions (Two-sided paired t test). Mechanical indentation evoked responses for units in a recording from an *Rorβ^iCre^; Vgat^FlpO^; R26^LSL-FSF-ReaChR-mCitrine^* mouse before and after NBQX application. Right: Summary from 3 *Rorβ^iCre^; Vgat^FlpO^; R26^LSL-FSF-ReaChR-mCitrine^* mice. Mechanically-evoked spikes were blocked in both Rorβ^+^ optotagged units (blue) and non-tagged units (grey; p < 0.01; Two-sided paired t test). **E.** Left: Mechanical responses to a series of step indentations for all optotagged inhibitory DH units (top) and average PSTH (bottom) from *Vgat^iCre^; R26^LSL-ChR2^* mice. Right: Assignment of optotagged Vgat^+^ interneurons to functional response clusters. **F.** As in (D), for simultaneously recorded putative excitatory DH interneurons (left), and cluster assignment (right). **G-K.** As in (D), for genetically distinct inhibitory interneurons (F-H): Rorβ (using *Rorβ^iCre^; Vgat^FlpO^*; *R26^LSL-FSF-ReaChR-mCitrine^* mice); PVi (*PV^2aCre^*; *Vgat^FlpO^*; *R26^LSL-FSF-ReaChR-mCitrine^*); Kcnip2 (*Kcnip2^CreER^; R26^ChR2-YFP^*); and excitatory interneurons (I-J): CCK (*CCK^iCre^; R26^ChR2-YFP^*) and PKCγ (*PKCγ^CreER^*; *R26^ChR2-YFP^* and *PKCγ^CreER^*; *Lbx1^FlpO^*; *R26^LSL-FSF-ReaChR-mCitrine^*). **L.** Spontaneous firing rates for all optotagged DH units. **M.** Truncated violin plots showing trough-to-peak durations for all optotagged units. **N.** Frequency- threshold tuning curves for genetically distinct interneuron populations.

**Figure S4.**
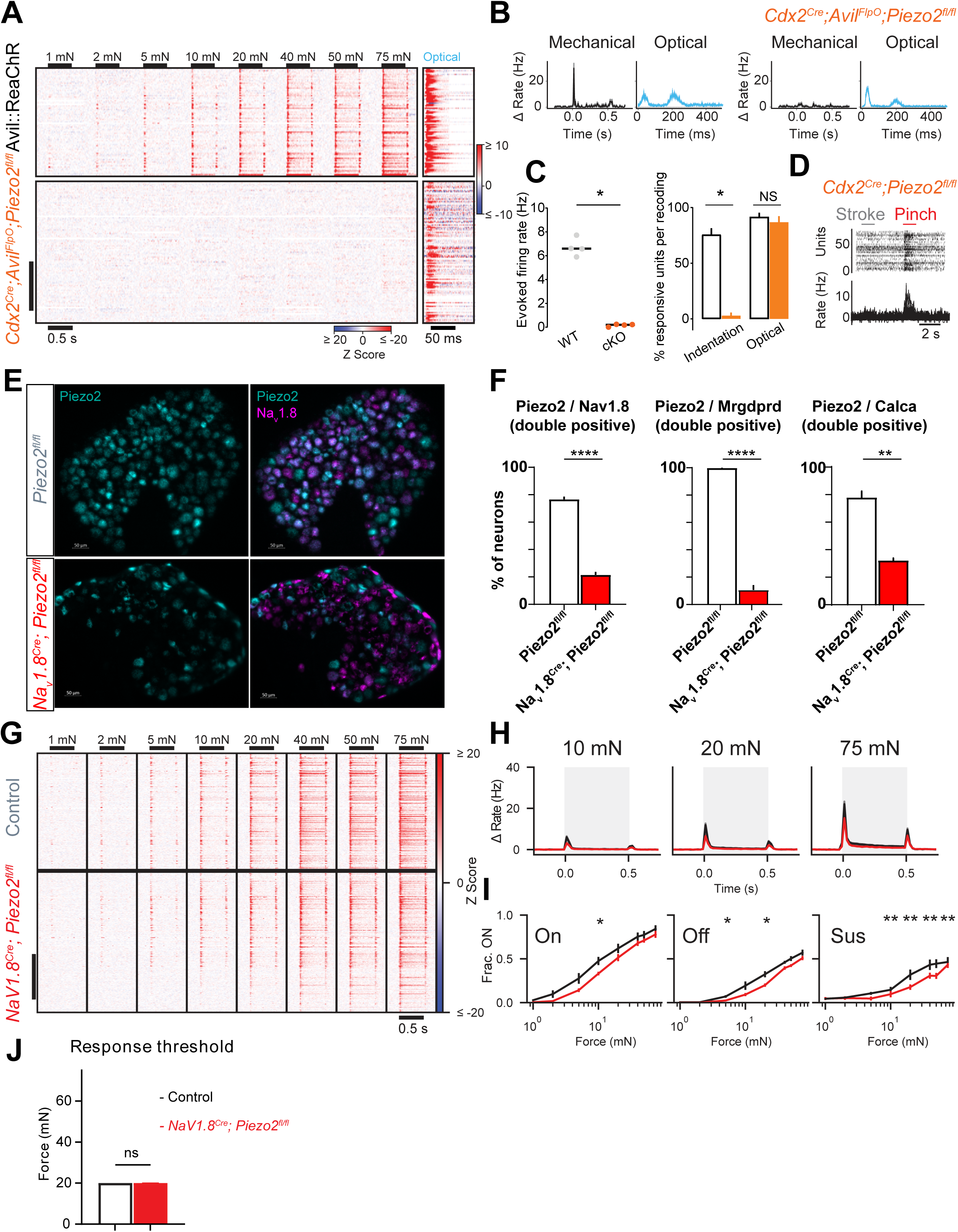
DH neuron responses in *Piezo2* conditional knockout animals. **A.** Mean firing rates in response to mechanical and optical activation of glabrous hindpaw in all DH units from littermate controls (*Piezo2^flox/flox^*; *Advillin^FlpO^* R26^LSL-FSF-ReaChR^; top; n= units, N= 3 mice) and *Cdx2-Cre; Piezo2^flox/flox^*; *Advillin^FlpO^* R26^LSL-FsF-ReaChR^ animals (bottom; N=3 mice). Units are sorted by depth. Note the timescale difference between mechanical and optical heatmaps. Scale bar: 50 units. **B.** Mechanical and optical responses to skin stimulation in example DH units from control (left) and *Cdx2-Cre; Piezo2^flox/flox^*; *Advillin^FlpO^* R26^LSL-FSF-ReaChR^ animals, showing both short and long-latency spikes to optical stimulation of the skin. **C.** Left: Summary of indentation-evoked firing rates in control (N=4) and *Cdx2-Cre; Piezo2^flox/flox^*; *Advillin^FlpO^* R26^LSL-FSF-ReaChR^ (N=2)/ *Cdx2-Cre; Piezo2^flox/flox^* (N=2) animals. Each dot represents one animal. Bars: mean. Right: Percentage of responding units to indentation and optical stimulation. Few units in Piezo2 mutants responded to indentation, however optical stimulation of the skin elicited similar amounts of responsivity in control and *Cdx2-Cre; Piezo2^flox/flox^*; *Advillin^FlpO^* R26^LSL-FSF-ReaChR^ animals. P < 0.05, Mann-Whitney U test. **D.** Example raster plot and PSTH of DH unit form Piezo2 cKO mouse showing no response to stroke on the skin, and robust response to noxious pinch. **E.** Representative images of Piezo2 and Na_v_1.8 smRNA-FISH from L1-L6 DRGs in *Na_v_1.8^Cre^; Piezo2^flox/flox^* animals (bottom), or *Piezo2^flox/flox^* littermates (top). **F.** Quantification of the percentage overlap between Piezo2 and DRG markers Na_v_1.8 (left), Mrgdprd (center), and CGRP (right). Multiple lumbar DRGs from three *Na_v_1.8^Cre^; Piezo2^flox/flox^* and three *Piezo2^flox/flox^*littermates were quantified. **E.** Z-scored firing rates of all dorsal horn units from littermate control (*Piezo2^flox/flox^*; top, N=3) and *Nav1.8-Cre; Piezo2^flox/flox^* (*Na_v_1.8* ^cKO^; bottom, N=3). Units are sorted by depth. Scale bar: 50 units. **F.** Grand mean baseline-subtracted firing rates (± SEM) to 10, 20 and 75mN step indentations across all units from control (black) and *Na_v_1.8* ^cKO^ (red) animals. **G.** Proportion of units from control (black) and *Na_v_1.8-Cre; Piezo2^flox/flox^* animals (red) responding at the ON (left), OFF (middle) and sustained (right) phase of step indentation across all forces tested. * p < 0.05, two-proportions Z test. **H.** Response thresholds from all units in control, and *Na_v_1.8-Cre; Piezo2^flox/flox^* animals. Bar: median with 95% CI. n.s., Mann-Whitney U test.

**Figure S5.**
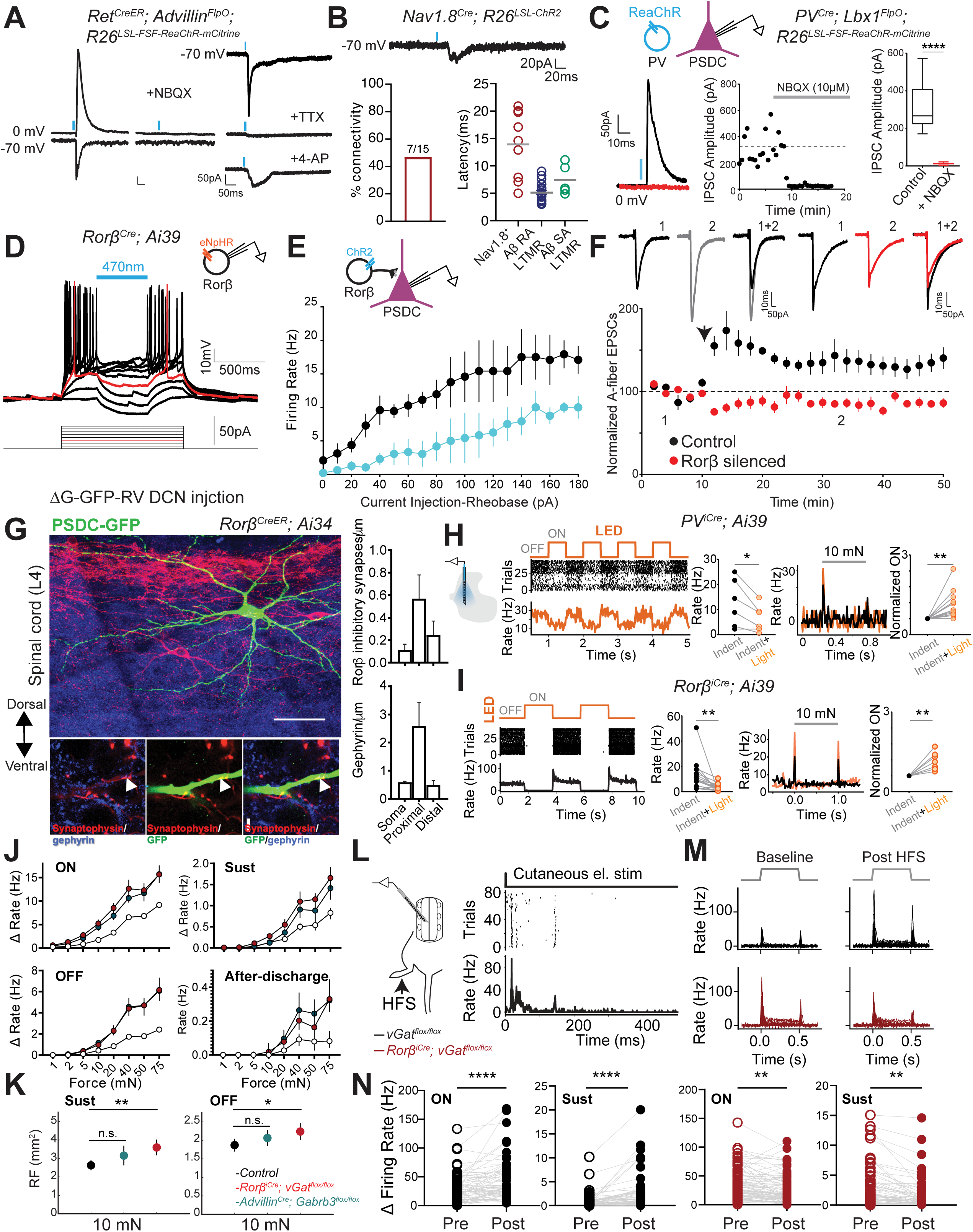
Sensory-evoked feed-forward and feedback inhibitory circuit motifs exert broad control over DH neuron responses. **A.** Optogenetic activation of Aβ-RA LTMR terminals evokes monosynaptic EPSCs and feedforward inhibition onto PSDC neurons. Left: Application of AMPA receptor antagonist NBQX (10uM) blocks both oEPSC and oIPSC. Right: Example EPSCs under baseline conditions and following co-application of voltage-gated Na+ channel blocker TTX and voltage- gated K+ channel blocker 4-aminopyridine (4AP). **B.** Top: Example EPSC in a PSDC neuron evoked by optogenetic activation of Nav1.8^+^ sensory terminals. Bottom: Connectivity rate for NaV1.8^+^ oEPSCs to PSDCs (left) and onset latencies for opto-evoked primary afferent EPSCs onto PSDC neurons (right). **C.** Top: Experimental design. Bottom: representative time course and example IPSC evoked by optogenetic activation of PV interneurons under baseline condition and following bath application of NBQX (left). Right: Average IPSC amplitudes before and after NBQX application (n=5 PSDCs; 3 mice). **D.** Example current clamp recording from eNpHR3.0-expressing Rorβ neuron. Light activation of eNpHR3.0 confirms silencing of spiking. Rheobase trace is shown in red. **E.** Summary current clamp recordings from PSDC neurons in response to current injection ramps paired with a 1s optical activation of Rorβ neurons (blue; n=6 neurons). **F.** Loss of LTP at Aβ-PSDC synapses in spinal cord slices form *Rorβ^iCre^; Cdx2FlpO; RC:PFtox* mice. Averaged EPSC amplitudes before and after high frequency dorsal root afferent stimulation (arrow) in PSDC neurons from control (n=3 PSDCs, 3 mice) and Rorβ silenced mice (n=3 PSDCs, 3 mice). Insets: Averaged EPSCs from single experiments before and 20 min after HFS. **G.** Right: Confocal images showing synaptophysin-tdTomato expression driven by *Ror*β*^CreER^*, and PSDC neurons (green) retrogradely labeled from DCN with glycoprotein (G) gene-deleted rabies virus (ΔG-GFP-RV). Co-labeling with gephyrin (blue) was used to determine axo-dendritic contacts from Rorβ^+^ inhibitory interneurons to PSDCs (arrowheads), quantified across the LTMR-RZ (right). **H.** *In vivo* MEA recording from *PV^iCre^*; R26^LSL-eNpHR3.0-YFP^ mice. Right: raster and PSTH for example PVi interneuron suppressed by eNpHR3.0-YFP, and quantification of the firing rate suppression in putative PVi interneurons (n=6 units; N=1 mouse). Left: example PSTH to 10mN step indentation without (black) and with (orange) acute PVi silencing (left), and quantification of the effect of PVi suppression of indentation-evoked spiking at the onset of step indentation in neighboring units on the probe (n=12 units). This effect was observed in 12 out of 48 units. **I.** *In vivo* MEA recording from *Rorβ^iCre^*; R26^LSL-eNpHR3.0-YFP^ mice. Left: Raster and PSTH for example Rorβ interneuron suppressed by eNpHR3.0-YFP, and quantification of firing rate suppression in all putative Rorβ interneurons (n=14 units; N=5 mice). Right: Example PSTH to 10mN step indentation without (black) and with (orange) acute Rorβ silencing, and quantification of the effect of Rorβ suppression on indentation evoked spiking at the ON response (n=7 units; N=5 mice) in neighboring units on the probe. This effect was observed in 7 out of 28 units. **J.** Median (±95% CI) firing rates for DH units in littermate control (white), FFI cKO (red) and PSI cKO (teal) mice in response to 1mN-75mM step indentations at the ON, OFF, sustained, and after-discharge periods. **K.** Average RF areas at 10mN force steps measured during the sustained (left) and OFF (right) phase of step indentations in control, FFI cKO and PSI cKO mice. **p < 0.01; *p < 0.05, standard deviation of bootstrapped RFs. **L.** Left: experimental design. Right: Raster plot and PSTH of DH unit in response to cutaneous electrical stimulation. Both short latency and long latency spikes were evoked, indicative of A and C fiber input. **M.** DH indentation responses to 75mN force steps before and after cutaneous HFS stimulation in control (black) and FFI cKO (red) mice. **N.** Summary firing rates evoked during ON and sustained phases of 75mN step indentations before and after cutaneous HFS stimulation in DH units from control (black), and FFI cKO mice. **p < 0.01; ****p < 0.0001, Mann-Whitney test.

**Figure S6.**
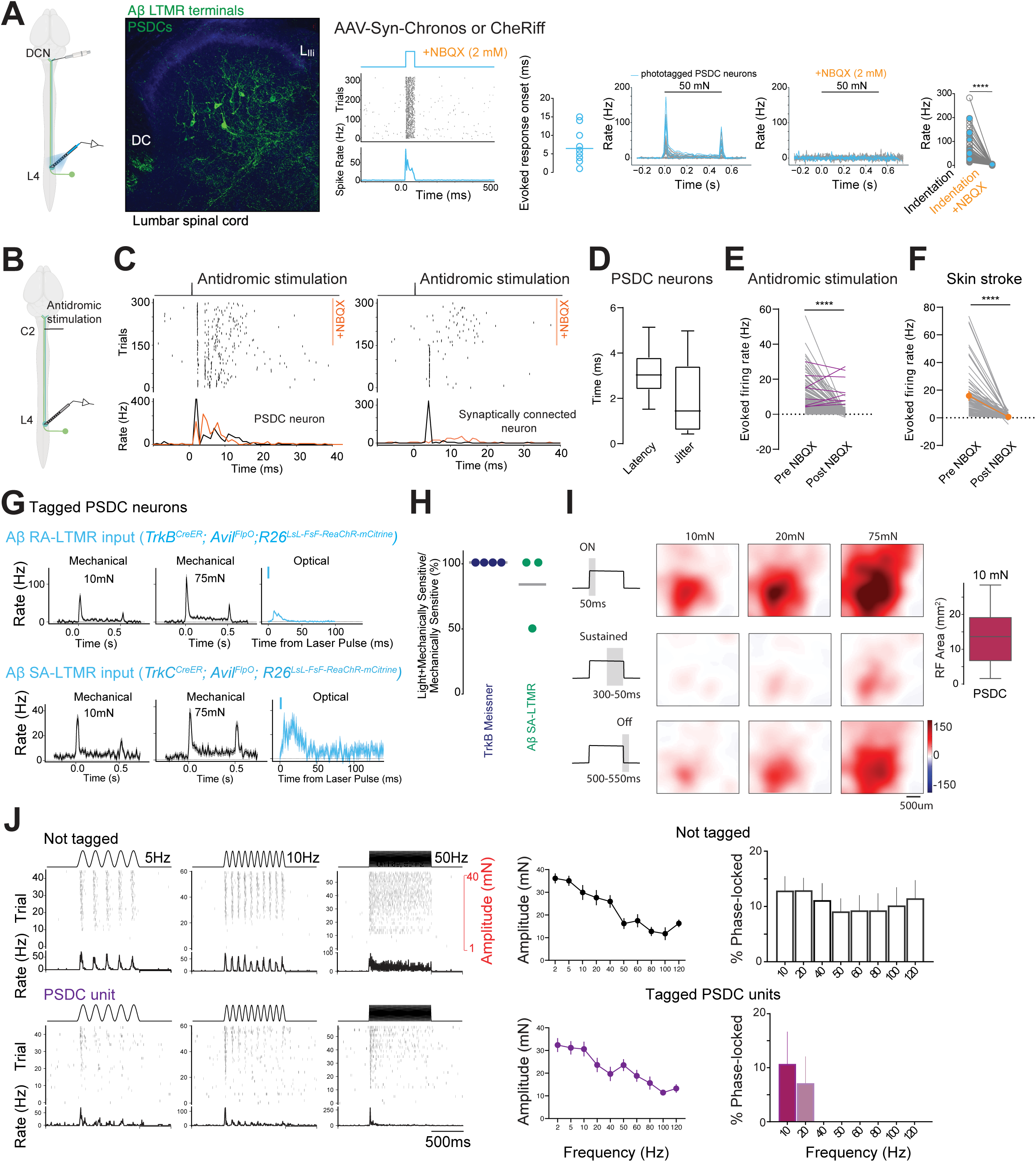
*In vivo* identification and mechanical response properties of PSDC neurons. A. Left: Schematic of PSDC retrograde viral labeling and recording configuration. This strategy results in excitatory opsin expression in both PSDCs and Aβ DCN projecting sensory neurons. Right: transverse lumbar spinal cord section with retrogradely labeled PSDCs and Aβ LTMR terminals (green), and IB4^+^ terminals (blue). B. Left: Example light-evoked raster plot and PSTH of PSDC neuron in the presence of NBQX (5mM) blocking excitatory synaptic transmission from direct pathway sensory neurons. Middle: summary light-evoked spike latencies in putative PSDC units. Right: Mechanically evoked responses for simultaneously recorded DH units before and after NBQX application to the surface of the cord. Mechanically-evoked spikes were blocked in both putative PSDC optotagged units (blue) and non-tagged units (grey; p < 0.01; Two-sided paired t test). C. Left: Dorsal column antidromic stimulation strategy. Right: Raster plot and PSTH in response to electrical antidromic stimulation of dorsal column axons at cervical levels C1-C3. Putative PSDC units continued to spike with short latencies and jitter in the presence of NBQX. D. Boxplots for all tagged PSDC neurons show low first spike latency and small jitter to DC antidromic stimulation. E. Summary of antidromic spike rates in putative PSDC neurons (purple) and neighboring units (grey) before and after NBXQ application. P < 0.0001; two-sided paired t test. F. Summary of firing rates in response to skin stroke before and after NBQX application confirms excitatory synaptic transmission blocking efficiency (mean: orange). P < 0.0001; two-sided paired t test. G. Tagged PSDC neurons receive direct input from Aβ-RA and Aβ-SA LTMRs. Mechanical and optically- evoked PSTHs (mean ± SEM) in example putative PSDC unit from *TrkB^CreER^*; *Advillin^FlpO^*; *R26^LSL-FSF-ReaChR-mCitrine^* (top) and from *TrkC^CreER^*; *Advillin^FlpO^*; *R26^LSL-FSF-ReaChR-mCitrine^* (bottom) mice. H. Proportion of mechanically sensitive PSDC units that respond to optical activation of distinct sensory neuron types in glabrous hindpaw. Each marker represents one animal. Bars: mean. I. Example PSDC spatiotemporal RF organization. J. Left: Example responses to mechanical vibrations in tagged PSDC neurons (bottom), and simultaneously recorded non-tagged DH units (top). Right: Phase-locked responses and vibration frequency tuning across all recorded PSDCs (purple) and simultaneously recorded nontagged units (black).

**Figure S7.**
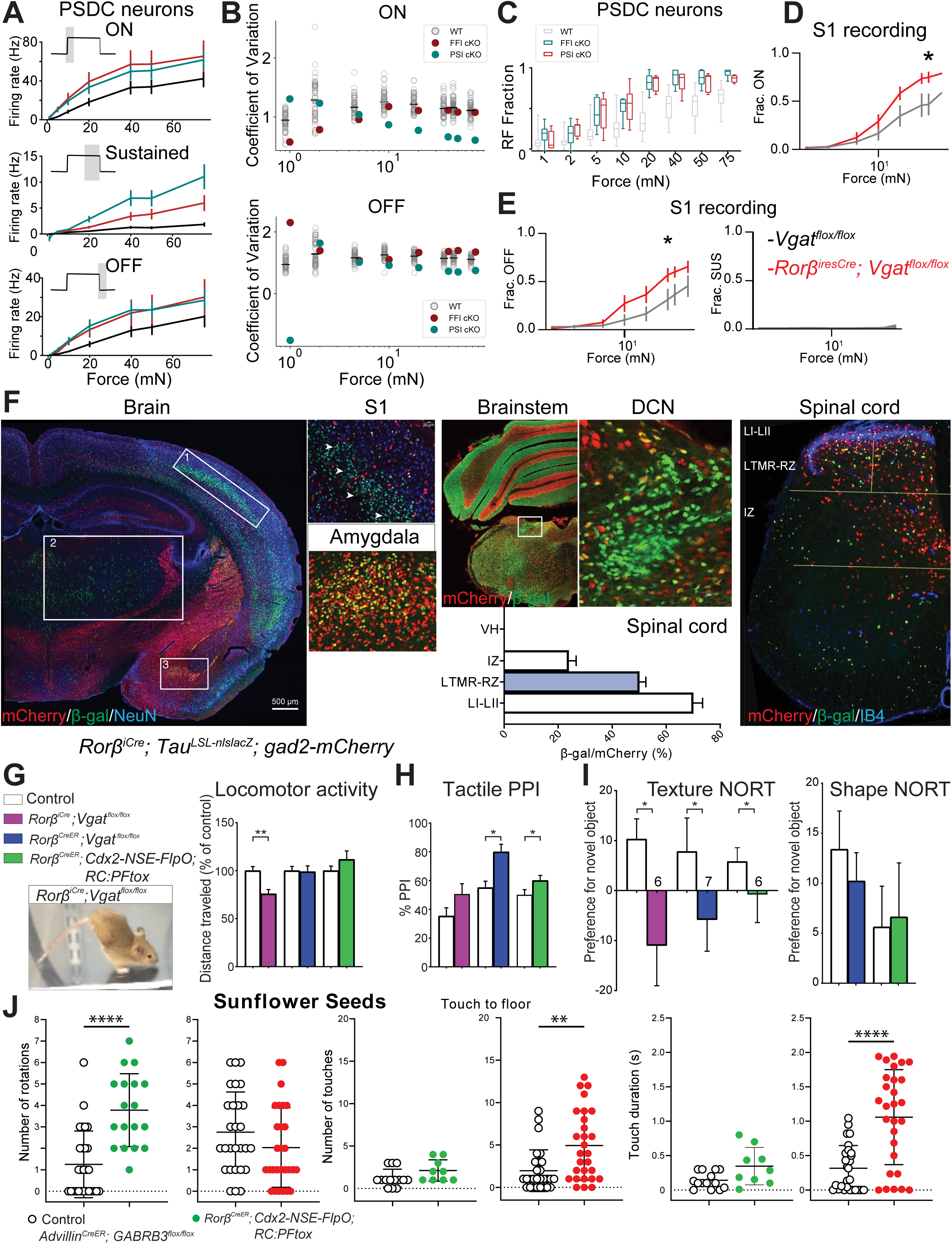
DH outputs shape tactile responses in S1 and somatosensory behaviors. **A.** Baseline-subtracted mean firing rates in tagged PSDC neurons from control, FFI cKO and PSI cKO mice at the ON, OFF, and the sustained portion of the indentation step. Error bars indicate SEM. **B.** C.V. of ON and OFF response magnitudes at indentation force steps between 2 to 75mN, measured from PSDC neurons in FFI cKO (red markers; n=17); PSI cKO (teal markers; n=14) and 50 representative sets of 14 randomly sampled PSDC neurons from control mice (grey markers). **C.** RF fraction of skin area mapped with increasing force steps in control PSDC neurons vs. PSDC neurons from FFI cKO and PSI cKO animals. **D-E.** Fraction (± SEM) of control and FFI cKO S1 units that produce a response at the ON (left), OFF (right) and sustained phase of the step indentation for each force. * p < 0.05; two-proportions Z test. **F.** Cross-sections through brain, brainstem and lumbar spinal cords from *Rorβ^iCre^; Tau^LSL-nlslacZ^; gad2-*mCherry animals (left to right). Colocalization of the inhibitory marker gad2 (red) with Rorβ lineage neurons (green) was found in lumbar spinal cord, but not at other sites of the somatosensory pathway including DCN, somatosensory thalamus or S1. Inhibitory Rorβ^+^ neurons were also found in amygdala and striatum. **G.** Genetic strategies used for selective silencing of spinal cord inhibitory Rorβ neurons. Right, silencing Rorβ neurons in *Rorβ^iCre^; Vgat^flox/flox^* causes characteristic hindlimb hyperflexion locomotor phenotype, previously described (Andre et al., 1998; Koch et al., 2017), demonstrating efficiency of the silencing strategy. **H.** Percentage inhibition of the startle response to a 125-dB noise, when the startle response was preceded by a light air puff (0.9PSI) in control littermates and Rorβ silenced mice. *p < 0.05. **I.** Discrimination indices for texture NORT and shape NORT in control littermates and Rorβ silenced mice. Positive value indicates preference for the novel object compared to the familiar object. *p < 0.05, Unpaired t-test. **J.** Sunflower seed assay reveals distinct sensory-motor phenotypes in DH FFI cKO (green) compared to PSI cKO (red) animals. Unpaired t test, **p < 0.01 and ****p < 0.0001.

## STAR METHODS

### Experimental model and subject details

All experimental procedures were approved by the Harvard Medical School Institutional Care and Use Committee (IACUC) and were performed in compliance with the Guide for Animal Care and Use of Laboratory Animals. Animals were housed in a temperature- and humidity-controlled facility and were maintained on a 12-hour light/dark cycle.

#### Mouse Lines

The following published mouse lines were used: *Vgat^iCre^* (JAX#016962; Vong et al., 2011), *Rorβ^iCre^* (JAX#023526; Harris et al., 2014), *Rorβ^CreER^* (JAX#030290; Abraira et al., 2017)*, Pvalb^2a-Cre^* (JAX#012358; Madisen et al., 2010), *Kcnip2-CreER* (JAX#030385; Abraira et al., 2017), *CCK^iCre^* (JAX#012706; Taniguchi et al., 2011), *PKC*γ*^CreER^* (JAX#030289; Abraira et al., 2017), *Lbx1^FlpO^* (Bourane et al., 2015), *Vgat-2A-FlpO* (JAX#029591; Daigle et al., 2018), *Gad2^T2A-NLS-mCherry^* (JAX#023140; Peron et al., 2015), Cdx2-Cre (Coutaud and Pilon, 2013), Cdx2-NSE-FlpO (JAX#030288; Abraira et al., 2017), *Advillin^Cre^* (Hasegawa et al., 2007), *Advillin^FlpO^* (JAX#032027; Choi et al., 2020), *TrkB^CreER^*(Rutlin et al., 2015), *Ret^CreER^* (Luo et al., 2009), *TrkC^CreER^*(Bai et al., 2015), *Na_v_1.8^Cre^* (JAX #036564; Nassar et al., 2004), *Calca^CreER^* (Song et al., 2012), *Calca-FlpE* (Choi et al., 2020); *Mrgprd^Cre^*(Rau et al., 2009), *Mrgprd^CreER^* (JAX#031286; Olson et al., 2017), *R26^LSL-FSF-ReaChR-mCitrine^* (JAX#024846; Hooks *et al*., 2015), *R26^LSL-ChR2-YFP^* (JAX#012569; Madisen et al., 2012), *R26^LSL-eNpHR3.0-YFP^* (JAX#014539; Madisen et al., 2012), *R26^LSL-synaptophysin-tdTomato^* (JAX#012570; Madisen et al., 2012), *Tau^LSL-mGFP-i-NLS-lacZ^* (JAX#021162; Hippenmeyer et al., 2005), *Vgat^flox^* (JAX#012897; Tong et al., 2008), *Gabrb3^flox^* (JAX#008310; Ferguson et al., 2007), *TrkB^flox^* (JAX#022362; Liu et al., 2012), *Atoh1^flox^* (JAX#008681; Shroyer et al., 2007), *Piezo2^flox^* (JAX #027720; Woo et al., 2014), *RC::PFtox* (Kim et al., 2009). Animals were maintained on mixed C57bl/J6, 129S1/SvImJ and CD1 backgrounds and included both males and females. C57Bl/J6 were obtained from Jackson Laboratories (000664).

**Table S1.** Genetic crosses and tamoxifen labeling strategy. Related to Figures 3,5 and 6.

| Strategy | Tamoxifen Age | Tamoxifen administration | Tamoxifen Dose (mg) |
| --- | --- | --- | --- |
| *Rorβ<sup>CreER</sup>*; *R26<sup>LSL-synaptophysin-tdTomato</sup>* | P21-P25 | intraperitoneal injection | 5mg (1mg/day) |
| *Kcnip<sup>CreER</sup>*; *R26<sup>LSL-ChR2</sup>* | P21-P25 | intraperitoneal injection | 5mg (1mg/day) |
| *PKCγ<sup>CreERT2</sup>*; *Lbx1<sup>FlpO</sup>*; *R26<sup>LSL-FSF-ReaChR</sup>*<br>*PKCγ<sup>CreERT2</sup>*; *R26* | P21-P25 | intraperitoneal injection | 5mg (1mg/day) |
| *Ret<sup>CreERT2</sup>*; *Advillin<sup>FlpO</sup>*; *R26<sup>LSL-FSF-ReaChR</sup>* | E10.5-E11.5 | Oral gavage | 3mg |
| *Ret<sup>CreERT2</sup>*; *R26<sup>LSL-FSF-ReaChR</sup>* | E10.5 | Oral gavage | 3mg |
| *TrkC<sup>CreERT2</sup>*; *Advillin<sup>FlpO</sup>*; *R26<sup>LSL-FSF-ReaChR</sup>* | E12.5 | Oral gavage | 3mg |
| *TrkB<sup>CreERT2</sup>*; *Advillin<sup>FlpO</sup>*; *R26<sup>LSL-FSF-ReaChR</sup>* | P3 | intraperitoneal injection | 0.5mg |
| *TrkB<sup>CreERT2</sup>*; *R26<sup>LSL-ReaChR</sup>* | P3 | intraperitoneal injection | 0.5mg |
| *CGRP<sup>CreER</sup>*; *R26<sup>LSL-ChR2</sup>* | P9 | intraperitoneal injection | 1mg |
| *Mrgprd<sup>CreERT2</sup>*; *R26<sup>LSL-ReaChR</sup>* | P9 | intraperitoneal injection | 1mg |

### Tamoxifen treatment

Tamoxifen was dissolved in ethanol (20mg/ml), mixed with an equal volume of sunflower seed oil (Sigma), vortexed for 5-10min and centrifuged under vacuum for 30min for ethanol removal. The solution was kept at -80°C and delivered via oral gavage to pregnant females for embryonic treatment (E10.5-E12.5, as specified in Table S1), or via intraperitoneal injection for postnatal treatment (P15-P25, as specified above). For spinal cord interneurons, time-points were chosen to label adult interneuron populations with defined anatomical and physiological signatures (Abraira et al., 2017). No changes in health or behavior were observed in tamoxifen treated animals compared to non-tamoxifen treated animals.

## METHOD DETAILS

### *In vivo* spinal cord multielectrode array electrophysiology

Adult (>6 weeks) animals were administered dexamethasone (2 mg/kg IP) 1-2 hours prior to recording to prevent tissue swelling, and were anesthetized with urethane (1.5 g/kg, Sigma). During surgery, isoflurane (1%) was administered but removed prior MEA recordings, and surgical plane of anesthesia was confirmed throughout the recording. The temperature of the animal was monitored and maintained (TC-344B, Warner Instruments) between 35 -37.5°C using a thermoelectric heater (C3200-6145, Honeywell) embedded in castable cement (Aremco). The hair surrounding the dorsal hump was shaved; a skin incision was made over the spinal segments T13 to L6, and the surrounding tissue was removed exposing the spinal column. An incision was made between vertebrae and tendons to allow for spinal clamp placement. The vertebrae above the recording site were stabilized using custom clamps to prevent movement. All tissue was cleared from vertebra and intervertebral space with forceps and spring scissors. The vertebrae between L4 and L5, were retracted to expose the dorsal surface of the spinal cord, the dura was removed and the surface of the cord bathed in mineral oil/ submerged in saline solution. A 32-channel silicon probe (Neuronexus A1x32-Poly3-5mm-25s-177-A32 or Cambridge Neurotech ASSY-37 H4 optrode) was inserted into the hindlimb representation region of the dorsal horn (medial L4-L5 spinal levels) and advanced up to ∼700 μm below the dorsal surface under visual guidance. Once positioned in a region where firing in many units could be evoked by brushing the hindpaw, the MEA was kept in place for 20 minutes to ensure a stable recording. Signals were amplified, filtered (0.1 - 7.5 kHz bandpass), and digitized (20 kHz) using a headstage amplifier and recording controller (Intan Technologies RHD2132 and Recording Controller). Data acquisition was controlled with open-source software (Intan Technologies Recording Controller version 2.07).

The hindpaw was stabilized with the plantar surface facing upwards, and stroking of the glabrous hindpaw was used a search stimulus to confirm probe placement. If cutaneous receptive fields were not on the glabrous hindpaw the probe was removed and reinserted in a new location. A 150-200 µm diameter Teflon-tipped indenter was controlled by a dual-mode force controller (Aurora Scientific 300C-I) and used to stimulate the glabrous hindpaw. For mapping receptive field (RF) areas, the position of the indenter was controlled with two linear translation piezo stages and a stage controller (Physik Instrumente U-521.24 and C-867.2U2). The position, force, and displacement of the indenter were commanded with custom-written Matlab scripts controlling a Nidaq board (National Instruments, NI USB 6259). Force steps were applied atop the minimum force required to keep the indenting probe in contact with the skin.

In a subset of experiments, electrodes were coated with DiI (Thermo Fisher) after the recording was completed, re-inserted in the spinal cord to the same coordinates and allowed to stabilize for 10-20 minutes. Animals were then anesthetized with isoflurane and transcardially perfused with PBS followed by 4% PFA for post hoc identification of the electrode track.

### Optical stimulation of genetically defined DH interneurons

To record from genetically defined DH interneuron populations, we used an optical tagging strategy in mice expressing excitatory opsins in specific DH neuron types (as specified in figure legends). We identified optotagged units by delivering 1-20ms pulses of blue light (4-10 mW/mm^2^ at fiber tip) to the surface of the spinal cord through optical fibers (200µm core diameter; NA = 0.66) attached to Cambridge Neurotech ASSY-37 H4 optrodes. Light was delivered from a 470 nm LED (M470F3, Thorlabs) or a 150 mW, 488 nm fiber coupled laser (OBIS LX #SKU 1220124, Coherent). Because most dorsal horn interneurons are glutamatergic, at the end of each experiment we applied ∼25-50µL NBQX (5 mM, Tocris, dissolved in H_2_O) to the surface of the cord to block recurrent excitation. Efficient block of glutamatergic synaptic transmission was determined by testing whether DH neuronal responses to brush stimulation or indentation were abolished, typically 10-20 minutes following NBQX application. Neurons that responded to optical stimulation both before and after NBQX application were considered optotagged. A modified stimulus-associated spike latency test (SALT; Kvitsiani et al., 2013; Emanuel et al., 2021) was additionally used to confirm short light-evoked spike latencies (<5ms) and low first spike jitter in optotagged units. We also verified the validity of optotagging by comparing the average peak-aligned sensory-evoked waveform with average light evoked waveform using Pearson’s correlation coefficient (r >0.9).

### *In vivo* dorsal column antidromic stimulation

In some experiments PSDC neurons were identified through antidromic activation of their dorsal column axons at cervical levels C1/C2. A laminectomy was performed to expose cervical spinal cord prior to lumbar dorsal horn recordings, and the region was sealed off with mineral oil.

After measuring tactile responses, a bipolar electrode (platinum-irridium 250 µm spacing, FHC) was placed on the surface of the dorsal column at cervical levels C1-C2. Single stimuli were applied at 0.1Hz frequency while evoked spikes were monitored in the lumbar spinal cord. This strategy activated both Aβ fiber axons and axons of PSDC neurons ascending via the dorsal column to the DCN. Therefore, evoked spikes could be measured in the majority of dorsal horn units that responded to low-threshold mechanical stimulation of the skin. To isolate antidromically activated PSDC units, ∼50µL NBQX (5 mM, Tocris, dissolved in H_2_O) was applied to the surface of the cord to block excitatory synaptic transmission from dorsal horn Aβ fiber synapses. Efficient block of glutamatergic synaptic transmission was confirmed by testing whether DH neuronal responses to brush stimulation or indentation were abolished, typically 10- 20 minutes following NBQX application. Units that responded to antidromic stimulation with reliable and precisely timed antidromic spikes (spike latencies and jitter <5ms) in the presence of NBQX were considered PSDC units.

### *In vivo* cortical recordings in awake mice

Primary somatosensory cortex (S1) MEA recordings were performed as described previously (Emanuel et al., 2021). Briefly, a craniotomy spanning hindpaw S1was performed at least 24h prior to the recording sessions (coordinates: 0.60 mm posterior and 1.65 mm lateral to bregma). For S1 MEA recordings of awake mice, a 32-channel silicon probe (Neuronexus A1x32-Poly2- 5mm-50s-177-OA32) was inserted into hindpaw S1 and the tip of the probe was advanced to 1100 mm below the dura.

Shortly after the probe was inserted into the brain, we confirmed probe placement in hindpaw S1 by gently brushing the skin of the animal with a fine paintbrush while monitoring spikes from multiple channels. If RFs were not on the glabrous paw, the probe was removed from the brain, moved to a new location within the craniotomy, and reinserted. Otherwise, the paw was tethered over a circular aperture (7.6 mm diameter) in an acrylic platform that supported the animal. A 0.5-mm diameter, cylindrical, Teflon-tipped indenting probe was controlled by a dual-mode force controller (Aurora Scientific 300C-I) and was used to stimulate the paw through the aperture. The position, force, and displacement of the indenter were commanded with the same custom Matlab (version 2017a) scripts controlling a Nidaq board (National Instruments, NI USB 6259) used for spinal cord recordings, described above. For multiregional RF mapping, we gently brushed the entire body of the animal. Regions of the body that elicited an increase in spiking to brushing were documented and included in the overall RF map.

### *In vivo* DRG electrophysiology

*In vivo* recordings were made from L4 DRGs using the same preparation as previously described (Bai et al., 2015; Neubarth et al., 2020; Emanuel et al., 2021) and a subset of the data presented here (Figure 5F and Figure S1C) originated from previously published recordings (Emanuel et al., 2021). Briefly, mice were treated with urethane (1 g/kg body weight) and anesthesia was supplemented with 1-2% isoflurane during the laminectomy. The L4 DRG was exposed via a dorsal incision and laminectomy. The exposed DRG was immersed in external solution containing (in mM) 140 NaCl, 3.1 KCl, 0.5 KH_2_PO_4_, 6 glucose, 1.2 CaCl_2,_ 1.2 MgSO_4_ (pH adjusted to 7.4 with NaOH) and the same solution was used to fill glass pipettes with a 20-30 µm tip diameter. Extracellular action potentials were measured using a Multiclamp 700A amplifier (Axon Instruments) operating in the voltage clamp configuration. The pipette voltage was set so that no current was flowing through the amplifier at baseline. The data was digitized at 40 kHz with a Digidata 1550a (Molecular Devices), low-pass filtered at 10 kHz (four-pole Bessel filter), and acquired using pClamp (Molecular Devices, Version 10). For MEA recordings, MEAs were inserted into the L4 DRG (H4-32ch, Cambridge Neurotech). MEA signals were high-pass filtered at 200 Hz, amplified (RHD2132) and acquired at 20kHz (RHD2000, Intantech) for offline processing.

### Spike sorting

Open-source software (JRCLUST version 3.2.2; Jun et al., 2017) was used to automatically sort action potentials into clusters, manually refine clusters, and classify clusters as single or multi units. Drift monitoring was performed during acquisition and experiments with detectable changes in spike waveforms were discarded. The voltage traces were filtered with a differentiation filter of order 3. Frequency outliers were removed with a threshold of 10 median absolute deviations (MADs). Action potentials were detected with a threshold of 4.5 times the standard deviation of the noise. Action potentials with similar times across sites were merged and then sorted into clusters with a density-based-clustering algorithm (Rodriguez and Laio, 2014) (clustering by fast search and find of density peaks) with cutoffs for log_10_(r) at -3 and log_10_(d) at 0.6. Clusters with a waveform correlation greater than 0.99 were automatically merged. Outlier spikes (> 6.5 MADs) were removed from each cluster.

To isolate putative single units, manual cluster curation was performed with JRCLUST split and merge tools. Clusters were classified as single units if *(1)* the waveforms were large with respect to baseline; *(2)* there was a clear refractory period in the cross-correlogram (interspike intervals > 1 ms); *(3)* waveforms were clearly distinct from nearby clusters. Spikes event times for clusters classified as single units were exported and processed in Python.

### Analysis of DH response properties and unbiased clustering Feature extraction and clustering

Automated unsupervised clustering was performed on DH single unit response profiles to 500ms step indentations of increasing intensities from 1mN to 75mN. For dataset collection, step indentations were applied either across the entire hindpaw (N= 106 mice; see Receptive field mapping below), or at manually determined RF centers for the majority of units on the probe (N= 36 mice). Units with spiking below the 1^st^ spiking percentile and units with no response threshold (for which only baseline firing was detected) were excluded from this analysis. The baseline firing rate was measured during the 1s interval before each indentation trial, and analysis was performed after the baseline firing rate was subtracted from stimulus-evoked responses.

We first extracted a minority of DH units inhibited by step indentations (i.e., units with ON and early sustained responses below their baseline firing rate at 40-75mN; n = 89). These neurons comprise the inhibitory functional cluster, cluster 6. We used principal component analysis (PCA) on the remaining dataset of 4971 units to extract features of indentation-evoked signals. The extracted features resembled classically defined temporal response profiles such as ON, OFF and sustained responses, with the first two principal components (PC1 and PC2) resembling RA- and SA-LTMR responses respectively (Figure S1D). We then performed k-medoid clustering using the number of PCs that account for 90% of variance in the data (n=503). Three different distance metrics (Euclidian, cosine and correlation distance) were tested, and cosine distance was chosen because this metric resulted in better cluster quality. To find the optimal number of clusters, silhouette values were calculated, and a local maximum determined (K= 5), with the addition of the inhibitory cluster, K=6 for the complete dataset. Peristimulus time histograms (PSTHs) with 10 ms time bins were generated to show the average response across all units within a functional cluster.

### Response feature analysis

Response properties of DH units across the six principal functional response types were then computed based on raw spike counts. Step indentation were subdivided into 5 distinct time windows to monitor different aspects of neuronal responses. The ON response was defined as the firing rate evoked during the first 50ms after the stimulus onset, Early Sustained response, as 50--200 msec following ON response; Late Sustained response, as 0-200 msec before stimulus offset; OFF response, as the first 50 msec after stimulus offset and after-discharge response as 50-200 msec after stimulus offset.

Thresholds for all units were determined by bootstrapping the baseline firing rate 1000 times to generate 95% confidence intervals, and detecting the smallest stimulus within the ON/OFF/Early Sustained or Late Sustained response windows that exceeded the upper bound or drops below the lower bound.

We quantified sensitivity and response magnitude by quantifying the maximum Z-scored firing rate within the 5 time intervals described above at all step indentations, and by determining the fraction of units that responded to each force step (Figures 4 and 5). A unit was determined to be responsive if it produced |z-scored firing rate| ≥ 3 between 10 and 50 ms after the onset or offset of the step indentation.

### Waveform characteristics

K-means clustering was performed using waveform statistics including trough-to-peak ratio, waveform slope, and trough-to-peak duration. S1 waveforms analyzed here originated from previously published recordings (Emanuel et al., 2021). K=3 was chosen to clearly separate waveforms in S1 into two regular spiking groups and one fast spiking group consistent with waveform clusters observed in mouse visual cortex (Jia et al., 2019). To compare spinal cord and S1 waveforms K=3 was used and revealed less separable groups in spinal cord waveforms.

### Mechanical and optical stimuli Step indentations

The amplitude of the ramp and hold indentation was 1mm, and their overall duration was 10 s, with on and off and ramps lasting 25 ms and separated by a 500 ms interval. Indentations were presented for a minimum of 30 trials at a unit’s hotspot (see below) or at multiple locations across the paw for receptive field mapping.

### Receptive field mapping

To measure glabrous receptive fields, step indentations of increasing forces were applied twice to randomized locations across the hind paw in a 6 x6 mm grid with 500 µm spacing between each stimulation site. In some cases, grid spacing was modified, as specified in the figure legends. The average grid area required to cover the entire glabrous hind paw was 36mm^2^. Receptive field area was calculated at forces delivered at and above each unit’s response threshold. To compute receptive fields, we first centered on each grid location and pooled all adjacent sites in a 3 x 3 grid into a larger spatial bin. For each response type, we then computed the bootstrap mean response over all 3 x 3 sites for 1000 bootstrap samples to establish 95% confidence interval. If the lower/upper bound of the CI was greater/smaller than the mean baseline firing rate of that bin, then an excitatory/inhibitory response was assigned to that center site. The excitatory/inhibitory receptive field size was then calculated as the fraction of sites that had an excitatory/inhibitory response multiplied by the probed area, whereas the RF fraction was simply the fraction of sites that had an excitatory/inhibitory response.

To measure responses at the receptive field hotspot, the average firing rate across all forces and all grid locations was computed, and grid locations with the largest responses (5% of the grid with the highest responsivity) were determined. The neuron’s hotspot was defined as the location on the skin that produced the highest firing rate when stimulated. 36 locations closest to the RF hotspot were selected, and responses across these sites were averaged, creating a PSTH, to represent responsivity at the hotspot. When response profile clustering was performed at the hotspot, only the data collected at hot spot grid locations was used.

To map receptive fields beyond glabrous hindpaw at locations where force-controlled step indentation delivery was not feasible, we used a hand-held brush head (5/0 Round Princeton Art & Brush Co., Blick) mounted to a strain gauge force sensor (MBL (BL341AH) 25 gram Model MBL load cell, Sensotec-Honeywell) connected to an amplifier (DMD-465WB, Omega). Stroke was delivered for 60 s to a given body region with 5 s inter-trial intervals. Mean baseline subtracted firing rate was measured for each body region and used to compute a preference index (related to Figure 7).

### Mechanical vibration stimuli and analysis

Vibratory stimuli were delivered to manually determined receptive field hotspots at intensities ranging from 1mN to 40mN and at 10 frequencies (2Hz to 120Hz). Frequency and amplitude of sinusoidal step vibrations, lasting 1s, were presented in a randomized order, separated by 1.5s interstimulus interval for a total of 250 trails. Units were determined to be vibrationally responsive if they fired action potentials at rates above baseline to at least two frequencies delivered at the 40mN intensity. The threshold for frequency tuned units is the lowest force evoking responses above baseline to three consecutive sine waves at each frequency.

Entrainment (phase-locking) was determined at intensities between 15mN and 40mN and at all frequencies. Entrained units responded with at least 0.5 spikes/cycle and displayed precise spike timing within a particular part of the sine wave. This was determined using a permutation test comparing actual spike times to randomized spike times.

### Optical skin stimulation and analysis

Pulses of light were generated every 100ms or 500ms using a 300mW, 445 nm laser (CST-H- 445-300, Ultralasers, Inc.). A minimum of 5000 light pulses were directed to the paw through two galvanometer scan mirrors (GVS201, Thorlabs) and an F-lens (FTH100-1064, Thorlabs), which focused the light to a 30 mm diameter spot. The intensity was modulated by inserting neutral density filters into the light path between the laser and the scan mirrors. Pulses were 1 ms in duration and the location of each pulse was randomized but confined to a 20 x 20 mm area that included the glabrous hind paw skin region. The location and timing of the light pulses were controlled using voltage signals generated with Matlab (2017b, Mathworks) and a National Instruments system (NI USB 6259). Z-scored firing rate was calculated in 1-ms bins using the baseline mean and standard deviation in the 10 ms preceding each laser pulse. Units were determined to be responsive to optical stimuli if the absolute value of the Z-scored firing rate exceeded 2.58 (98% confidence interval) within 25 ms after the laser pulse for A-fiber activation, or within 200 ms for C-fiber activation.

### Optical RF measurements

Optical RFs for DH and DRG neurons were computed with 1mm^2^ spatial bins. Baseline-subtracted optically-evoked firing rate across these 1mm^2^ subregions was normalized to the maximum optically-evoked firing rate. Bins with responses > 0.5 of the maximum-normalized response were included in the overall RF area for each unit (calculated as the sum of the binarized subregions). DRG optical RFs originated were analyzed from previously published recordings (Emanuel et al., 2021).

### PSDC retrograde labeling

Animals (P13-15 for slice physiology experiments; 4-6weeks for *in vivo* electrophysiology; P21- P30 for histology) were anesthetized with isoflurane and placed in a stereotaxic frame. The head was tilted 30° forward. Puralube ointment was applied to the eyes. The hair over the neck and caudal scalp was removed using a clipper, and the skin was sanitized using betadine. An incision was made in the midline of the back skin at the cervical level to expose neck muscles and local anesthetic (0.5% lidocaine) was applied to the incision site. Neck muscles were removed to expose the brainstem. A small incision was made on the dura to expose the DCN. The following retrograde tracers were injected into the DCN using a glass pipette under visual guidance: Adeno-Associated Virus (AAV9- hSyn-CheRiff-TdT, titer 2.73E+15 in 0.9% saline, Boston Children’s viral core; AAV2retro-hSyn-Chronos-GFP; Boston Children’s viral core titer 1.3 E+13 in 0.9% saline, Boston Children’s viral core), Rabies Virus (RabV-deltaG-GFP, titer 5.84E+7 - 9.48E+8 IU/mL, Boston Children’s viral core), or cholera toxin subunit B (CTB; 2 mg/ml in PBS, Invitrogen). Once penetrating the surface, a small volume (30-50 nL) of tracer was injected at multiple locations in the DCN (200-400 nl total volume). The pipette was then removed, and overlying muscle and skin was stitched together with sutures. Animals were administered analgesic (Buprenex SR, 0.1 mg/kg) and monitored post-operatively. At the appropriate time point (4 weeks following AAV injections or 3-7 days following CTB or RabV injections), mice were used for electrophysiology experiments or transcardially perfused for tissue harvest.

### Spinal cord slice preparation

Mice were briefly anesthetized with isoflurane, and intracardially perfused with ice-cold oxygenated choline solution (ACSF) prior to spinal cord removal. The isolated spinal cord was embedded in low-melting agarose (Sigma Aldrich), and transverse slices (300 μm) with dorsal roots attached were prepared from lumbar levels (L3-L5) using a Leica vibrating blade microtome (Leica VT1200S). Spinal cord slices were prepared in ice-cold oxygenated choline solution containing (in mM): 92 Choline Chloride, 2.5 KCL, 1.2 NaH_2_PO_4_, 30 NaHCO_3_, 20 HEPES, 2.5 Glucose, 5 Sodium Ascorbate, 2 Thiourea, 3 Sodium Pyruvate, 10 MgSO_4_ 7H_2_O, 0.5mM CaCl_2_ 2H2O. Slices recovered at 34°C for 30min in HEPES holding solution equilibrated with 95% O_2_, 5% CO_2_ containing (in mM): 86 NaCl, 2.5 KCl, 1.2 NaH_2_PO_4_, 35 NaHCO_3_, 20 HEPES, 25 glucose, 5 NaAscorbate, 2 Thio Urea, 3 Na Pyruvate, 1 MgSO_4_ 7H_2_O, 2 CaCl_2_ (pH 7.3, osmolarity 305; Ting et al., 2014), and were held in the same HEPES solution at room temperature until use.

### Acute slice recordings

Spinal cord slices were transferred to a submerged recording chamber at room temperature and continuously perfused with ACSF containing (in mM): 2.5 CaCl_2_, 1 NaH_2_PO_4_, 119 NaCl, 2.5 KCl, 1.3 MgSO_4_ 7H_2_O, 26 NaHCO_3_, 25 dextrose, and 1.3 Na ascorbate, saturated with 95% O_2_, 5% CO_2_ at a rate of ∼1-2 ml/min. Cells were visualized using infrared differential interference contrast and fluorescence microscopy. Whole cell voltage-clamp recordings of retrogradely labeled PSDCs in laminae IV-V were obtained under visual guidance using a 40x objective.

Whole cell voltage clamp recordings were obtained using an internal solution containing (in mM): 135 CsMeSO_3_, 4 ATP-Mg^2+^, 0.3 GTP-Na^+^, 1 EGTA, 3.3 QX-314(Cl^-^ salt), 8 Na_2_-Phoshocreatine and 10 HEPES. Synaptic currents were evoked with electrical stimulation of dorsal roots using a suction electrode at Aβ fiber strength (<=25 mA, 20-100µs; Nakatsuka et al., 2000; Torsney and MacDermott, 2006). For isolation of sensory-evoked EPSCs and feedforward IPSCs, PSDC neurons were voltage-clamped alternatively at the reversal potential for synaptic inhibition (-70mV) and excitation (0mV). To activate ChR2 in acute slices, LED whole field illumination was used through a water immersion 40x objective. Aβ-LTMR axon terminals were stimulated with brief pulses (1-5ms) of blue light (473 nm, 5mW). Optically-evoked IPSCs (oIPSCs) were blocked by inhibitory or excitatory transmission blockers as specified in the figure legends.

For plasticity experiments, sensory-evoked glutamatergic EPSCs in PSDCs were isolated using pharmacological antagonists of GABA_A_Rs and glycine receptors (10 µM bicuculline, 1 µM strychnine). High-frequency stimulation (HFS; two 1 s trains at 100 Hz, intertrain interval 20 s, at 1.5 times test current intensity) was delivered after a stable 10 min baseline.

Current-clamp recordings were obtained with an internal solution containing (in mM): 135 K-Gluconate, 10 NaCl, 2 MgCl_2_, 0.5 EGTA, 10 HEPES, 2 Mg-ATP, 0.3 Na-GTP. Input resistance and access resistance were monitored continuously throughout each experiment and cells were excluded from analysis if these values changed by more than 10% during the experiment or if the resting membrane potential was higher than 50 mV. Data were acquired using a Multiclamp 700B amplifier, a Digidata 1440A acquisition system, and pClamp 10 software (Molecular Devices). Sampling rate was 10 kHz, and data were low-pass filtered at 3 kHz. No correction for junction potential was applied.

### Spinal cord immunohistochemistry of free-floating sections

Mice (P30-P35) were anesthetized with CO_2_ and perfused with 5-10mL modified Ames Media (Sigma) in 1x PBS, followed by 20-40 mL of 4% paraformaldehyde (PFA) in PBS at room temperature (RT). Vertebral columns (including spinal cords and dorsal root ganglia) were dissected and were post-fixed in 4% PFA at 4°C for 2-16 hr. Lumbar spinal cord sagittal sections (100-150 µm) were cut on a vibrating blade microtome (Leica VT100S) and processed for immunohistochemistry as described previously (Hughes et al., 2012; Abraira et al., 2017).

Briefly, tissue samples were rinsed in 50% ethanol/water solution for 30 min to allow for enhanced antibody penetration. Three washes in high salt Phosphate Buffer Saline (HS PBS) were conducted each lasting 10 min. The tissue was then incubated in primary antibodies in high salt Phosphate Buffer Saline containing 0.3% Triton X-100 (HS PBSt) for 48-72 hr at 4°C. Primary antibodies used goat anti-mCherry (1:1000, AB0040, Sicgen), rabbit anti-GFP (1:1000, A-11122, Thermo Fisher Scientific) and mouse anti-gephyrin (1:1000; Synaptic Systems). The tissue was washed in HS PBSt, then incubated in a secondary antibody solution in HS PBSt overnight at 4°C. Secondary antibodies included species-specific Alexa Fluor 405, 488, 546, and 647 conjugated IgGs (Life Technologies). Tissue sections were then mounted on glass slides, coverslipped with Fluoromount Aqueous Mounting Medium (Sigma) and stored at 4°C.

### Immunohistochemistry of frozen tissue sections

Brains and vertebral columns, including spinal cords and dorsal root ganglia, were removed from perfused mice and post-fixed in 4% PFA at 4 **°**C overnight. Tissues were washed in 1× PBS for over 3 h, and brains and spinal cords were finely dissected out from the rest of the tissue. Brain and spinal cord tissues were cryoprotected in 30% sucrose at 4 **°**C for 2 days, embedded in OCT (1437365, Fisher), frozen using dry ice and stored at –80 **°**C. Coronal brain sections and transverse spinal cord sections (30–40 μm) were cryosectioned on a cryostat (Leica). Spinal cord sections were collected on glass slides (12-550-15, Fisher), and brain sections were collected on glass slides or in 1× PBS. Sections were washed three times for 5 min with 1× PBS containing 0.1% Triton X-100 (0.1% PBST), incubated with blocking solutions (0.1% PBST containing 5% normal goat serum (S-1000, Vector Labs) or normal donkey serum (005-000-121, Jackson ImmunoResearch) for 1h at RT, incubated with primary antibodies diluted in blocking solutions at 4 **°**C overnight, washed three times for 10 min each with 0.1% PBST, incubated with secondary antibodies diluted in blocking solutions at 4 **°**C overnight, washed again four times for 10 min each with 0.1% PBST and mounted with Fluoromount-G (Southern Biotech). For spinal cord sections, IB4 (1:500; Alexa 647 conjugated, L21411, Molecular Probes) was incubated together with secondary antibodies. Primary antibodies used include goat anti-mCherry (1:1,000, AB0040, Sicgen), chicken anti-GFP (1:1,000, GFP-1020, Aves Labs), rabbit anti-GFP (1:1,000, A-11122, Thermo Fisher Scientific), mouse anti-NeuN (1:1,000, MAB377, Millipore).

Secondary antibodies included Alexa 488–conjugated donkey anti-chicken antibodies, Alexa 546–conjugated donkey anti-goat antibodies and Alexa 647– conjugated donkey anti-mouse antibodies. All secondary antibodies were purchased from Thermo Fisher Scientific and Jackson ImmunoResearch Labs, and used at 1:500 dilution.

### DH synaptic connectivity analysis

Synaptic connectivity analysis between Rorβ and PSDC neurons was performed in *Ror*β*^CreER^; R26^LSL-synaptophysin-tdTomato^* mice in which PSDC neurons were retrogradely labeled with dG-RV-GFP (see above), as previously described (Abraira et al. 2017). In addition to the genetically encoded presynaptic marker synaptophysin, the inhibitory postsynaptic marker gephyrin was used in this analysis. A total of 4 animals were used.

Z stack images of spinal cord slices were taken on a Zeiss LSM 700 confocal microscope using a 40X oil-immersion lens (Zeiss; Plan-Apochromat 40X/NA 1.40) and scanned at a z-separation of 0.5 µm. Images were taken in DH lamina IIiv-IV, which was defined as between the lamina IIiv border (marked by IB4 binding) and 250 µm below that border. Imaging parameters (laser power, gain/offset, averaging, dwell time, etc.) were consistent across animals. For analysis, images were first pre-processed using ImageJ: using the channel of PSDC labeling, two masks were generated – one using a standardized threshold for signal in this channel and a second by expanding this first mask by 1 µm in all dimensions. These masks were then used to isolate pre- and post-synaptic labeling by multiplying these channels (using the Image Calculator function) with the expanded and non-expanded masks, respectively. Next, images were analyzed using the Cell Counter ImageJ plugin to determine the proportion of inhibitory reporter terminals that apposed a gephyrin-immunoreactive puncta. Puncta were counted as a function of location: cell body, proximal neurite (within the first 50 µm) or distal neurite (beyond the first 50 µm).

### In situ hybridization

Detection of Piezo2, Na_V_1.8, CGRP and *Mrgprd* transcripts was performed by fluorescent in situ hybridization, as previously described (Lehnert *et al*., 2021; Sharma et al., 2020). Briefly, lumbar DRG ganglia were rapidly dissected from euthanized mice, frozen in dry-ice and stored at -80°C until further processing. DRGs were cryosectioned at a thickness of 20 μm and RNA was detected using RNAscope (Advanced Cell Diagnostics) according to the manufacturer’s protocol. The following probes were used: Mm *Piezo2* exons 43-45 (Cat# 439971-C3), *Na_V_1.8* (Cat#: 426011-C2), *Calca* (Cat#:420361), *Mm-Mrgprd* (Cat#: 417921). Sections were mounted in FluoroMount-G (Fisher 0100-01) and imaged on a Zeiss LSM 700 confocal microscope using a 10x or 20x objective.

### Behavior

#### Prepulse Inhibition (PPI) Assay

Tactile PPI Assay was used as a measurement of hairy skin sensitivity, as described previously (Orefice et al., 2016). Briefly, using a San Diego Instruments startle reflex system (SR-LAB Startle Response System), mice were tested in a cylindrical chamber within a soundproof chamber. For tactile PPI, a prepulse of an air puff was administered to the hairy skin of the back. The air puff was delivered at a constant intensity (0.9 PSI), at varying intervals before the startle pulse, ranging from 50ms to 1s. A tone pre-stimulus (ranging from 68 dB to 80 dB, for 20ms), followed by startle tone stimulus (120 dB, 20 ms) version of the PPI assay (acoustic PPI) was done as a control. Acoustic PPI was done with background noise set at 65dB, while tactile PPI had background noise at 75dB to ensure the air puff prepulse could not be heard. Startle reflex was quantitated using an accelerometer measuring the amplitude of movement of the animal.

#### Texture NORT

Texture NORT was used to measure texture discrimination in glabrous skin, as previously described (Orefice et al., 2016). On day 1 and 2 of behavioral testing, animals were individually habituated to an empty testing chamber (40 cm x 40 cm x 40 cm) under dim lighting for 10 min. Following habituation, animals were tested on color/shape NORT (day 3) and texture NORT (day 4). During the learning phase, the animal explored the testing chamber, in which two identical objects spaced equidistant from each other and the chamber walls, for 10 min. The animal was returned to the home cage for a 5 min retention period, during which the chamber and objects were thoroughly cleaned with 70% ethanol and one of the objects was replaced with a novel object. After the 5 min period, the animal was returned to the chamber for the 10 min exploration of the testing phase. Both the learning and testing phases were video-recorded from above. Custom MATLAB scripts were used to track animal position in the chamber and calculate the amount of time the animal spent investigating the objects. For color/shape NORT, the objects were wooden blocks differing in shape and color. In texture NORT, the objects were plexiglass cubes (4 cm^3^) that were visually identical but varied in texture (rough or smooth).

Animals were whisker plucked three days before the start of habituation. The preference for the novel object was calculated based on the time exploring both objects.

#### Sunflower Seed Assay

To test the role of feed-forward and presynaptic inhibition in a fine sensorimotor behavior, we used a Sunflower Seed Handling Assay, as described previously (Neubarth et al., 2020). To habituate animals, one week prior to testing one to two tablespoons of black oil sunflower seeds (Bio-Serv, S5137-1, Wagner’s, 76025) were added to the floor of the animals’ home cage for five consecutive days. If animals did not recognize seeds as a food source, then a teaspoon of seeds were cracked before adding to the cage floor.

Habituation to the behavior chamber and handling began two days prior to sunflower seed testing. Animals were habituated to the behavior room environment and investigator handling by undergoing tail inking on habituation day 1. To ink the tail, each animal was gently lifted and placed on the cage wire food hopper facing away from the investigator. Firmly holding the tail midway from the tail base, a blue permanent soy ink marker was rolled across the tail forming parallel lines to indicate identifying ear notch numbers. Once inked animals were gently transferred back into the home cage to await test chamber habituation.

The test chamber was constructed of a black matte acrylic wall and three optically clear walls, 10 in (l) x 8 in (w) x 8 in (h), 0.25 in thick, centered on a white matte acrylic floor under diffuse warm white light (2700K). Three digital USB 2.0 CMOS video cameras mounted on camera sliders were positioned on each clear side of the test chambers. One additional overview camera was mount directly above the test chamber.

Two days prior to testing and during testing seeds were withheld from the home cage to encourage foraging and seed eating in the test chamber. Animals were not food restricted for this assay. On habituation day 1 and day 2, animals were removed from the home cage and placed in an empty test chamber resting on the white matte acrylic floor. Each animal was given 2-3 seeds while freely exploring the test chamber for 20 minutes. Following the exploration animals were then returned to their home cages. Testing began on day 3. Animals were transferred from their home cage and placed in the test chamber and allowed to explore the chamber for 5 minutes.

Following acclimation, 2-3 seeds were placed on the floor of the test chamber and seed eating activity was recorded. At the completion of the seed eating test animals were removed from the test chamber and returned to their home cage. Chambers were reset and cleaned with unscented soapy water, wiped down with ddH2O and dried. Animals that failed to eat seeds after 20 minutes were returned to their home cage and the test rescheduled. This schedule was repeated until each animal fully deshelled and consumed multiple seeds. Seeds that were partially deshelled/consumed or discarded were not counted.

Behaviors were measured by defined seed deshelling and eating actions: (i) Seed peeling and deshelling—the act of grasping and holding the sunflower seed between the forepaws, clamping the upper and lower incisors into the shell surface, and applying downward force (dip) that pushed the shell away from the head and teeth toward the floor. This action resulted in a systematic peeling of the shell to expose the seed kernel. Animals unable to maintain a firm grip on, or fully grasp, the shell would typically adapt by touching, tapping, resting and/or bracing the seed against the floor between the forepaws. Animals also “tucked” the shell against their abdomen, holding the shell between the forepaws, clamping their upper and lower incisors onto the shell surface pulling their heads backward away from the shell and forepaws to peel off sections of the shell exposing the seed kernel. (ii) Touch-taps—the act of touching and/or holding and/or bracing the seed shell between the forepaws and the floor during seed peeling. (iii) Dip—the act of holding the seed shell between forepaws, clamping shell between incisors, and applying downward force to peel off sections of shell. (iv) Rotate—the act of or ability to change and/or manipulate shell orientation within the forepaws. (v) Rocking—the act of grasping the shell between the forepaws with the shell firmly between incisors, using forepaws to “rock” the shell side to side between the incisors to bite into the shell to peel and expose the seed kernel.

#### Quantification and statistical analysis

Statistical tests were conducted using the SciPy stats module (Python 3.8.5) or GraphPad Prism. Both non-parametric tests and parametric tests were used, depending on data normality, for comparing two independent groups (Mann-Whitney U test or Student’s t test), and multiple groups (Kruskal-Wallis H test/ one-way ANOVA or two-way ANOVA for multiple groups with multiple timepoints). All statistically tests performed are indicated in the figure legends or supplemental statistics table. p < 0.05 was considered significant. Additional details on sample sizes and statistical tests for each experiment can be found in the figure legends, main text, and the supplemental statistics table S1.

All data reported in this study will be shared by the lead contact upon request. Code is available upon request to the corresponding authors.

